# Hierarchical control of remyelination by heparan sulfate sulfation

**DOI:** 10.64898/2026.09.24.754157

**Authors:** Rupadevi Muthaiah, Roopa Ravichandar, Rathnakar Poddeti, Aneri P. Shah, Bhargavi Kulkarni, Trina Rudra, Ding Xu, Nagendra K Rai, Ranjan Dutta, Fraser J. Sim

## Abstract

Efficient myelin regeneration requires coordinated signaling between oligodendrocyte progenitor cells (OPCs) and the extracellular matrix, yet how specific heparan sulfate sulfation patterns regulate this process remains poorly understood. The extracellular endosulfatases Sulf1 and Sulf2 selectively remove 6-*O*-sulfate groups from heparan sulfate proteoglycans (HSPGs). We previously demonstrated that these enzymes are highly expressed by OPCs following demyelination and inhibit OPC recruitment, differentiation, and remyelination through activation of BMP and WNT signaling. We investigated whether sulfotransferases that establish 2-*O* and 6-*O* sulfation during HSPG biosynthesis regulate remyelination. Using OPC-specific conditional knockout models, deletion of the 2-*O* sulfotransferase *Hs2st1* enhanced OPC recruitment, oligodendrocyte differentiation, and remyelination through suppression of BMP and WNT signaling. Loss of 2-*O* sulfation induced compensatory remodeling of the heparan sulfate glycome, with a marked enrichment of 6-*O*-sulfated HS species. Conversely, deletion of 6-*O* sulfotransferases impaired OPC recruitment and differentiation and reversed the *Hs2st1* phenotype, identifying 6-*O* sulfation as the dominant functional determinant of remyelination. Together, these findings establish hierarchical control of remyelination by heparan sulfate sulfation, demonstrating that compensatory remodeling of the heparan sulfate glycome governs regenerative signaling following demyelination.

## INTRODUCTION

The microenvironment within regions of demyelination profoundly influences the recruitment, survival, and differentiation of oligodendrocyte progenitor cells (OPCs) that are essential for efficient myelin repair (Ghorbani and Yong, 2021). The extracellular matrix (ECM) comprises approximately 20% of brain volume and serves to coordinate signaling cues regulating cellular growth and synaptogenesis. Chondroitin sulfate proteoglycans (CSPG) and hyaluronan are well-established components of the inhibitory extracellular milieu that impairs remyelination (Sobel and Ahmed, 2001; Back et al., 2005) and are dynamically regulated during OPC differentiation (Mironova et al., 2026). Demyelinated lesions in multiple sclerosis (MS) are characterized by the deposition of a diverse ECM, including heparan sulfate proteoglycans (HSPGs) (van Horssen et al., 2005). Expression of HSPG genes are upregulated in MS lesions (Mohan et al., 2010) and following induction of experimental demyelination (Macchi et al., 2020).

HSPGs consist of a core protein decorated with one or more covalently attached heparan sulfate (HS) glycosaminoglycan chains composed of repeating disaccharide units of glucuronic acid (GlcA) and *N*-acetylglucosamine (GlcNAc). During biosynthesis in the Golgi apparatus, these chains undergo extensive and highly regulated modifications catalyzed by specific enzymes, including *N*-deacetylation, *N*-sulfation (*Ndst1-4*), C5 epimerization of GlcA to iduronic acid (*Glce*), and *O*-sulfation at 2-*O* (*Hs2st1*), 6-*O* (*Hs6st1-3*), or 3-*O* (*Hs3st1-6*) positions. Following their presentation at the cell surface, HS chains are further dynamically remodeled by extracellular endosulfatases (*Sulf1/2*), which selectively remove 6-*O* sulfate groups, and by heparanase, which cleaves the HS backbone (Esko and Selleck, 2002; Sarrazin et al., 2011).

Together, these biosynthetic and post-synthetic modifications generate structurally distinct HS domains that confer ligand specificity and fine-tune downstream signaling pathways (Esko and Selleck, 2002; Sugahara and Kitagawa, 2002; Lin, 2004). Consistent with the “sugar code” hypothesis, discrete HS sulfation patterns uniquely regulate signaling events critical for nervous system development and repair (Allen and Rapraeger, 2003; Lee and Chien, 2004; Holt and Dickson, 2005; Kreuger et al., 2006; Carlsson and Kjellen, 2012). We have previously demonstrated that *Sulf1/2* act as inhibitors of OPC differentiation following demyelination (Saraswat et al., 2021b). Pharmacological and genetic blockade of sulfatase activity following spinal cord demyelination accelerated oligodendrogenesis and remyelination through attenuation of BMP- and WNT-mediated signaling. Furthermore, in an IFN-γ induced model of inflammatory demyelination, heparanase activity contributed to impaired OPC recruitment and axonal damage (Saraswat et al., 2021a). Together, these findings highlight HS sulfation dynamics on OPCs and other cell types as critical regulators of signaling pathways that govern myelin repair and its failure.

During development, OPCs in mice robustly express a relatively restricted repertoire of sulfotransferases, including *Hs2st1*, *Hs6st1*, and *Hs6st2* (Zhang et al., 2014). In this study, we investigated how OPC-intrinsic sulfotransferase activity shapes the extracellular signaling environment following demyelination. We hypothesized that selective modulation of HS sulfation on OPC-expressed HSPGs influences the efficiency of remyelination. Loss of 2-*O* sulfation was dispensable for repair and instead triggered a compensatory increase in 6-*O* sulfation that was itself necessary to promote oligodendrogenesis and enhance remyelination.

## MATERIALS AND METHODS

### Animals and surgery

All experiments were performed according to protocols approved by the University at Buffalo’s Institutional Animal Care and Use Committee. *NG2CreER; Rosa26YFP* animals were a gift from Akiko Nishiyama (University of Connecticut, Storrs, CT) (Zhu et al., 2011). *NG2CreER; Hs2st1^fl/fl^*mice were generated by crossing *Hs2st1^fl/fl^* mice to *NG2CreER* transgenic mice (Stanford et al., 2010). Sperm from *Hs6st1^fl/fl^*(Izvolsky et al., 2008) mice were provided by Dr. Jeffrey Esko’s lab. The sperm was used to perform *in vitro* fertilization on *NG2CreER; Hs2st1^fl/fl^*mice (Roswell Park Comprehensive Cancer Center). The female progeny were subsequently subjected to *in vitro* fertilization with sperm from *Hs6st2^−/−^*mice (Nagai et al., 2013). This led to the generation of five distinct conditional HS sulfotransferase transgenic colonies: (1) *NG2CreER; Hs2st1^fl/fl^,* (2) *NG2CreER; Hs6st1^fl/fl^,* (3) *NG2CreER; Hs6st2^−/−^,* (4) *NG2CreER; Hs6st1^fl/fl^; Hs6st2^−/−^,* and (5) *NG2CreER; Hs2st1^fl/fl^; Hs6st1^fl/fl^; Hs6st2^−/−^*. Cre-mediated recombination was induced by intraperitoneal administration of tamoxifen (200 mg/kg, Sigma) every other day for a total of five injections, with the final injection administered seven days prior to surgery. All experimental mice received tamoxifen in an identical manner, and littermate controls lacking Cre were compared with Cre-expressing conditional knockout (cKO) mice.

Focal demyelination in the spinal cords of young adult mice, aged 8-11 weeks, was induced using established protocols (Welliver et al., 2018; Ravichandar et al., 2024). In this procedure, the mice were first anesthetized with isoflurane, and 0.5 μL of 1% lysolecithin (Lα-lysophosphatidylcholine, Sigma) was injected into both the dorsal and ventral funiculi of the spinal cord. This injection was performed between two adjacent thoracolumbar vertebrae to facilitate targeted demyelination and study its effects on spinal cord functionality. Post-operative analgesia was provided by subcutaneous injection of Ethiqa XR (3.25 mg/kg). Terminal 5-ethynyl-2’-deoxyuridine (EdU) was administered by IP injection at 48 h and 24 h prior to euthanasia (50 mg/kg).

### Human MS tissue

MS tissue was prepared as described (Izvolsky et al., 2008; Tripathi et al., 2019). Briefly, brains were collected as part of the tissue procurement program approved by the Cleveland Clinic Institutional Review Board. Informed consent was obtained from all tissue donors. Tissue specimens were not considered “human subjects” due to the absence of interaction with living patients and the use of autopsy materials under HHS regulations 45 CFR Part 46. Brains were removed according to a rapid autopsy protocol. Chronic active demyelinated lesions were analyzed from six secondary progressive MS patients (2M, 4F) and one primary progressive MS patient (1F) (52-70 years old, disease duration >5 years, postmortem interval <9 h).

### Animal tissue processing and analysis

Animals were euthanized at 3-, 5-, 7-, or 14-day post-lesion (dpl) via transcardial perfusion with saline followed by 4% paraformaldehyde under deep anesthesia. Tissue processing and identification of lesions were conducted as previously described (Ravichandar et al., 2024). Slides adjacent to the lesion centers, identified by solochrome cyanine staining, were used for all immunohistochemical procedures. Sections were permeabilized for 30 min with 1% Triton X-100 (Alfa Aesar, Ward Hill, MA) and 0.25% Tween 20 (Calbiochem, San Diego, CA). Following permeabilization, sections were blocked for 1 h with a solution containing 0.5% Triton X-100 and 5% normal serum (goat serum unless otherwise indicated) (Thermo Fisher Scientific, Waltham, MA). The primary antibodies used included human anti-HS20 (1:200), mouse anti-10E4 (Amsbio, 370255-S, 0.1 µg/ml), rabbit anti-Olig2 (Millipore, AB9610, 1:500), mouse anti-CC1 (Millipore, OP80, 1:50), mouse anti-GFAP (Sigma, G3893, 1:300), rabbit anti-Iba1 (Wako Chemicals USA, 019-19741, 1:300), cleaved caspase-3 (Asp175) (Cell Signaling, 9661S, 1:250), mouse pan axonal neurofilament (BioLegend, SMI-312R, 1:1000), mouse pan neuronal neurofilament (BioLegend, SMI-311R, 1:1000), rabbit β-Amyloid Polyclonal (APP1) (Thermo Fisher Scientific, 512700, 1:250), and Timp1 (Abcam, ab216432, 1:200). Alexa 488, 594, and 647 conjugated secondary antibodies (Invitrogen, Carlsbad, CA) were applied at a dilution of 1:500. To validate the specificity of immunostaining for cell surface sulfated motifs, tissue sections were treated with or without 25 mU/ml heparin lyase III (HEP III) in a reaction buffer composed of 25 mM HEPES, 150 mM NaCl, and 2 mM CaCl₂ (pH 7.0) at 37 °C for 2 h. Following enzymatic treatment, sections were blocked in 5% normal serum in 1X phosphate-buffered saline (PBS) for 1 h. Primary and secondary antibodies were diluted in the same blocking solution. Images of spinal cord sections were acquired at 20× magnification using widefield epifluorescence (Olympus IX83). For each marker, the average signal intensity was quantified from at least two sections per animal. Lesions with a cross-sectional area smaller than 10,000 µm² or those extending into adjacent gray matter were excluded from analysis. All quantifications were performed by an investigator blinded to the sample identities.

### Mouse and human OPC isolation and immunocytochemistry

For isolation of human primary OPCs, fetal brain tissue samples, between 17 and 22 weeks of gestational age, were obtained from Advanced Bioscience Resources (Alameda, CA). Informed consent was obtained from all donors. Following review by the University at Buffalo Research Subjects Institutional Review Board, the tissue acquisition and research were determined not to involve human subjects, as defined under HHS regulations 45 CFR 46.102(f). Forebrain samples were minced and dissociated using papain and DNase as previously described (Sim et al., 2011). Magnetic sorting of CD140a/PDGFαR^+^ cells was performed as described (Conway et al., 2012). Human OPCs (hOPCs) were maintained on plates coated with poly-ornithine and laminin, in neural differentiation (ND) media supplemented with 20 ng/mL PDGF-AA (PeproTech, Cranbury, NJ) and 5 ng/mL NT-3 (PeproTech), as described (Abiraman et al., 2015). Mouse OPCs (mOPCs) were isolated from *NG2CreER; Hs2st1^fl/fl^*(Cre^+^) and corresponding wildtype (WT) littermates following injection of 4-hydroxytamoxifen (4 consecutive days, IP, 100 mg/kg). The brains for *Hs2st1* cKO and WT littermates were dissociated as above, and MACS was performed using anti-mouse CD140a beads (Miltenyi Biotec). The mOPCs isolated were cultured at 5 x 10^4^ cells/mL in 6-well plates coated with poly-ornithine and laminin in ND media with PDGF-AA (10 ng/mL) and FGF (10 ng/mL, Peprotech). Two-thirds of the culture media was replaced every other day.

The expression of sulfotransferases involved in 2-*O* sulfation and the distribution of sulfated HS motifs were assessed using Hs2st1 and 10E4 antibodies. Mouse OPCs were maintained as progenitors, fixed, and immunostained with mouse anti-Hs2st1 (Santa Cruz, sc-376530, 1:50) and mouse anti-10E4 (Amsbio, 370255-S, 1:50), followed by co-labeling with rabbit anti-Olig2 (Millipore, AB9610MI, 1:400). For analyzing HS and *N*-sulfated HS expression in human OPCs, the cells were similarly maintained as progenitors, fixed, and stained with mouse anti-3G10 (Amsbio, 370260-S, 1:50) and mouse anti-10E4, along with co-labeling using rabbit anti-Olig2 (Millipore, AB9610MI, 1:400). The 3G10 antibody detects a neoepitope generated by HEP III digestion of HS. Therefore, prior to staining, the cells were treated with or without 25 mU/ml HEP III in a reaction buffer containing 25 mM HEPES, 150 mM NaCl, and 2 mM CaCl₂ (pH 7.0) at 37 °C for 2 h. Following enzymatic treatment, cells were blocked in 5% normal serum (from the host species of the secondary antibody) in 1X PBS for 1 h. After overnight incubation with primary antibodies, Alexa Fluor 488- and 594-conjugated secondary antibodies (Invitrogen) were applied at 1:500 dilutions. Images were captured at 20× magnification using widefield epifluorescence microscopy (Olympus IX83). The specificity of both 10E4 and 3G10 antibodies was independently validated by HEP III treatment, which abolished 10E4 immunoreactivity and unmasked the 3G10 neo-epitope following HS chain cleavage (**Supplementary Fig. 1**), thereby confirming selective detection of intact versus cleaved HS structures.

### RNA extraction and Quantitative RT-PCR

Total RNA was isolated from demyelinating white matter lesions (WML) in six chronic progressive MS brain samples. Normal-appearing white matter (NAWM) tissue surrounding the lesion was used as a control. Total RNA was isolated with E.Z.N.A Total RNA Kit (Omega Bio-tek, 74104) and reverse transcribed with random hexamers to cDNA with SuperScript VILO cDNA Synthesis Kit (Invitrogen, 11754050), according to manufacturers’ instructions. The expression of the HS-related genes: HS2ST1 (Hs00202138_m1, Applied Biosystems, Foster City, CA), HS6ST1 (Hs00757137_m1), HS6ST2 (Hs00708234_s1), SULF1 (Hs00392834_m1), and SULF2 (Hs01016480_m1) was assayed by TaqMan-based qPCR (4453320). Gene expression was normalized to GAPDH (Hs02786634_g1). Each sample was run in triplicate. ΔCt values were used to determine relative expression changes using the ΔΔCt method (2^-ΔΔCt^). Samples were compared using Student’s *t-*tests, *p* <0.05 was considered statistically significant.

### Western blot

For cellular *Hs2st1* expression analysis, mOPCs were washed in PBS, lysed, and sonicated in a buffer containing 50 mM β-glycerophosphate, 20 mM HEPES, 1% Triton X-100, protease inhibitors (Roche, Mannheim, Germany), phosphatase inhibitor cocktail 2 (Sigma-Aldrich, St.Louis, MO, USA), and phosphatase inhibitor cocktail 3 (Sigma-Aldrich). Protein concentrations were determined using a Bradford assay (Bio-Rad, Hercules, California, USA), and 30 µg protein was loaded onto 10% polyacrylamide gels for Western blot. Proteins were transferred to nitrocellulose membranes and blocked in PBS containing 5% milk and 0.1% Tween-20 (Calbiochem, San Diego, CA, USA) in PBS. Primary antibodies against Hs2st1 (Thermo Fisher Scientific, PA5-89472, 1:2000), and β-actin (Cell Signaling Technology, 3700, 1:10,000) were applied overnight at 4 °C in blocking buffer. HRP-conjugated secondary antibodies (Sigma-Aldrich) were diluted 1:5000 in blocking buffer. Blots were imaged using the ChemiDoc MP imaging system (Bio-Rad).

### Electron microscopy

For assessment of remyelination, tissue was processed as described in Welliver et al. (2018). Briefly, mice were euthanized at 14 dpl by transcardial perfusion with 2% glutaraldehyde in 0.1 M phosphate buffer, and spinal cords were extracted (*n* = 3-6 per group). One millimeter thick blocks surrounding the spinal cord lesion were post-fixed in osmium tetroxide (EMS, Hatfield, PA, USA), dehydrated through ascending ethanol washes, and embedded in TAAB resin (TAAB Laboratories, Aldermaston, UK). One-micrometer sections were cut, stained with 1% (w/v) toluidine blue (Sigma-Aldrich, St.Louis, MO, USA), and examined by light microscopy to identify lesions. Selected blocks with lesions were trimmed, ultrathin sections cut, and examined by transmission electron microscopy (Hitachi, HT7800). Images were acquired at 2,500x magnification. Analyses of remyelinated axons and g-ratios were performed blinded. For remyelination counts, a minimum of 800 axons was counted for each animal from at least six different fields, with all animals included per treatment group. Analysis of g-ratio (axon diameter/fiber diameter) was performed as described in Dillenburg et al. (2018). Briefly, axon and fiber diameters were measured using FIJI (diameter = 2 × √[area/*π*]). A minimum of 100 axons was analyzed per animal. The frequency distribution of axon diameter and *g*-ratio of remyelinated axons was calculated by binning on axon diameter and g-ratio.

### BaseScope and RNAScope In situ hybridization

BaseScope™ was performed to distinguish WT and *Hs2st1* KO mRNA transcripts. To detect WT *Hs2st1*, we used a 1ZZ probe (NPR-0049759; BA-Mm-Hs2st1-E1E2) targeting nucleotides 456-493 of NM_001355234.1. This region is absent in the KO transcript due to a deletion spanning nucleotides 193-483 (partial exon 1). For detection of the *Hs2st1* KO transcript, a separate 1ZZ probe (NPR-0049758; BA-Mm-Hs2st1-cE1E2) was designed to target nucleotides 168-492 of NM_001355234.1, incorporating the 193-483 nt deletion. This probe spans a cryptic exon junction generated by the deletion and does not hybridize to the WT transcript. For the BaseScope assay, fixed sections were baked at 60 °C for 1 h, then washed with ethanol, and subjected to tissue pretreatment. The sections were exposed to probes spanning BA-Mm-Hs2st1-E1E2 and BA-Mm-Hs2st1-cE1E2 for 2 h in the ACD HybEZ Hybridization System. BaseScope Detection Reagents (323800; Advanced Cell Diagnostics Bio, Newark, CA) AMP 0-AMP 12, FastRed and FastGreen were applied according to manufacturer instructions and washed using wash buffer. To visualize nuclei, sections were counterstained in Gill’s Hematoxylin I. Positive signals of WT mRNA of *Hs2st1* were identified as blue punctate dots present in the nucleus and cytoplasm, and positive signals of KO mRNA of *Hs2st1* were identified as red punctate dots present in the nucleus and cytoplasm. All tissues were tested using negative control probes to control for nonspecific binding. For all BaseScope-based imaging, brightfield images of spinal cords were acquired (Olympus IX51).

RNAScope was performed with probes targeting *Pdgfra1* (GenBank: NM_011058.2), *Apcdd1* (GenBank: NM_133037.3), and *Id4* (GenBank: NM_031166.2) mRNA by using the RNAScope Fluorescent Multiplex Detection Kit (Cat. No. 323100; Advanced Cell Diagnostics Bio) according to the manufacturer’s instructions. Dissected mouse spinal cords were cryopreserved in sucrose (7.5 and 15% w/v) for 24 h each and were snap frozen in OCT and sectioned coronally at 16 µm thickness. Fixed sections were baked at 60 °C for 1 h, washed with ethanol, followed by tissue pretreatment, and probe hybridization according to the RNAScope multiplex protocol. Sections were counterstained with DAPI to visualize nuclei. Positive signals were identified as punctate dots present in the nucleus and nearby cytoplasm. Negative control probes were included to control for nonspecific binding. Images were acquired at 20x magnification using widefield epifluorescence microscopy (Olympus IX83).

### Extraction of HS from the cells

HS extraction and disaccharide analysis were performed as previously described, with minor modifications (Wang et al., 2020; Wang et al., 2022). Briefly, cell pellets were suspended in water and proteolyzed with pronase E (10 mg per 1 g protein, w/w) at 55 °C for 24 h. The digest was denatured at 100 °C for 10 min and centrifuged at 14000 rpm for 10 min, and the supernatant was collected. A ^13^C-labeled N-sulfo heparosan recovery calibrant was added to the supernatant prior to DEAE column purification (Wang et al., 2020).

DEAE column mobile phase A consisted of 20 mM Tris (pH 7.5) and 50 mM NaCl, and mobile phase B consisted of 20 mM Tris (pH 7.5) and 1 M NaCl. After sample loading, the column was washed with 1.5 mL mobile phase A, followed by elution of the HS fraction with 1.5 mL mobile phase B. The eluate was desalted using a YM-3 kDa molecular weight cutoff spin column, and the retentate subjected to digestion with heparin lyases. Heparin lyase (HEP) digestion was performed in a total volume of 100 μL containing 7.5 μL of enzymatic buffer (100 mM sodium acetate and 2 mM calcium acetate buffer, pH 7.0, with 0.1 g/L BSA), 4 μL HEP I (0.7 mg/mL), and 2.5 μL HEP II (13.6 mg/mL). The reaction mixture was incubated at 37 °C overnight. Prior to recovery of the digestion products, a known amount of ^13^C-labeled disaccharide calibrants was added. HS disaccharides were recovered by centrifugation, and the collected supernatant was freeze-dried prior to AMAC derivatization. Eight ^13^C-labeled HS disaccharide calibrants were used for quantitative analysis of non-3-*O*-sulfated HS species, as previously described (Wang et al., 2020).

### Chemical derivatization of HS disaccharides

AMAC (2-Aminoacridone) derivatization of lyophilized HS disaccharides was performed as previously described (Wang et al., 2020). Briefly, samples were resuspended in 5 μL of 0.1 M AMAC in DMSO/glacial acetic acid (17:3, v/v) and incubated at room temperature for 15 min. Subsequently, 5 μL of freshly prepared 1M aqueous sodium cyanoborohydride was added, and the reaction mixture was incubated at 45 °C for 2 h. Following incubation, samples were centrifuged, and the supernatant was collected for LC-MS/MS analysis.

### LC-MS/MS analysis

AMAC-labeled HS disaccharides were analyzed using a Vanquish Flex UHPLC System (Thermo Fisher Scientific) coupled to a TSQ Quantis triple-quadrupole mass spectrometer (Wang et al., 2020). Separation was achieved on a C18 column (Agilent InfinityLab Poroshell 120 EC-C18, 2.7 μm, 4.6 × 50 mm). Mobile phase A consisted of 50 mM ammonium acetate in water, and mobile phase B consisted of methanol. The elution gradient was programmed from 5 to 45% mobile phase B over 10 min, followed by isocratic 100% mobile phase B for 4 min and then re-equilibration in isocratic 5% mobile phase B for 6 min, at a flow rate of 0.3 mL/min.

Detection was performed using online triple-quadrupole mass spectrometry operating in the multiple-reaction-monitoring (MRM) mode. Electrospray ionization was conducted in negative-ion mode with the following parameters: spray voltage 4.0 kV, sheath gas 45 Arb, auxiliary gas 15 Arb, ion transfer tube temperature 320 °C, and vaporizer temperature 350 °C. Data acquisition and processing were performed using TraceFinder software.

### Statistical Analysis

All quantification and data analysis were performed by an investigator blinded to sample identity. Statistical analyses were performed using GraphPad Prism (GraphPad Software, San Diego, CA). Data were compared using Student’s *t* test, one-way ANOVA, or two-way ANOVA, as appropriate, and *p* < 0.05 was considered statistically significant. Representative micrographs are shown from experiments performed with at least three biologically independent replicates. For experiments including more than three biologically independent replicates, the number of replicates is provided in the corresponding figure legends. Data are presented as mean ± SEM.

## RESULTS

### HS 2-*O* and 6-*O* sulfation is dynamically upregulated following demyelination and enriched in MS lesions

To define how heparan sulfate (HS) sulfation changes during demyelination and repair, we used focal lysolecithin-induced demyelination in the adult mouse spinal cord, a model with well-characterized temporal kinetics of OPC recruitment, differentiation, and remyelination (**Fig. 1a**) (Fancy et al., 2011). In young adult mice, OPC recruitment peaks around 5 days post-lesion (dpl), differentiation initiates by 7 dpl, and remyelination is evident by 14 dpl, enabling a detailed assessment of extracellular matrix remodeling during repair. Highly sulfated HS epitopes associated with 2-*O* and 6-*O* sulfation were detected using HS20 antibody (Gao et al., 2016), and antibody specificity confirmed by pretreatment with heparinase III, which abolished immunoreactivity (**Fig. 1b**). Quantitative analysis of HS20 revealed a pronounced increase in sulfated HS within demyelinated lesions over time (**Fig. 1c-e**). Mean fluorescence intensity (MFI) was significantly elevated at 7 dpl and further increased at 14 dpl compared to earlier time points (3 and 5 dpl; *n* = 3-4 mice per group; one-way ANOVA, F(4,12) = 17.5, p < 0.0001) (**Fig. 1e**). Notably, this increase coincided temporally with OPC differentiation and active remyelination rather than initial progenitor recruitment, suggesting that HS 2-*O* and 6-*O* sulfation is selectively enriched during later stages of lesion repair.

**Figure 1:**
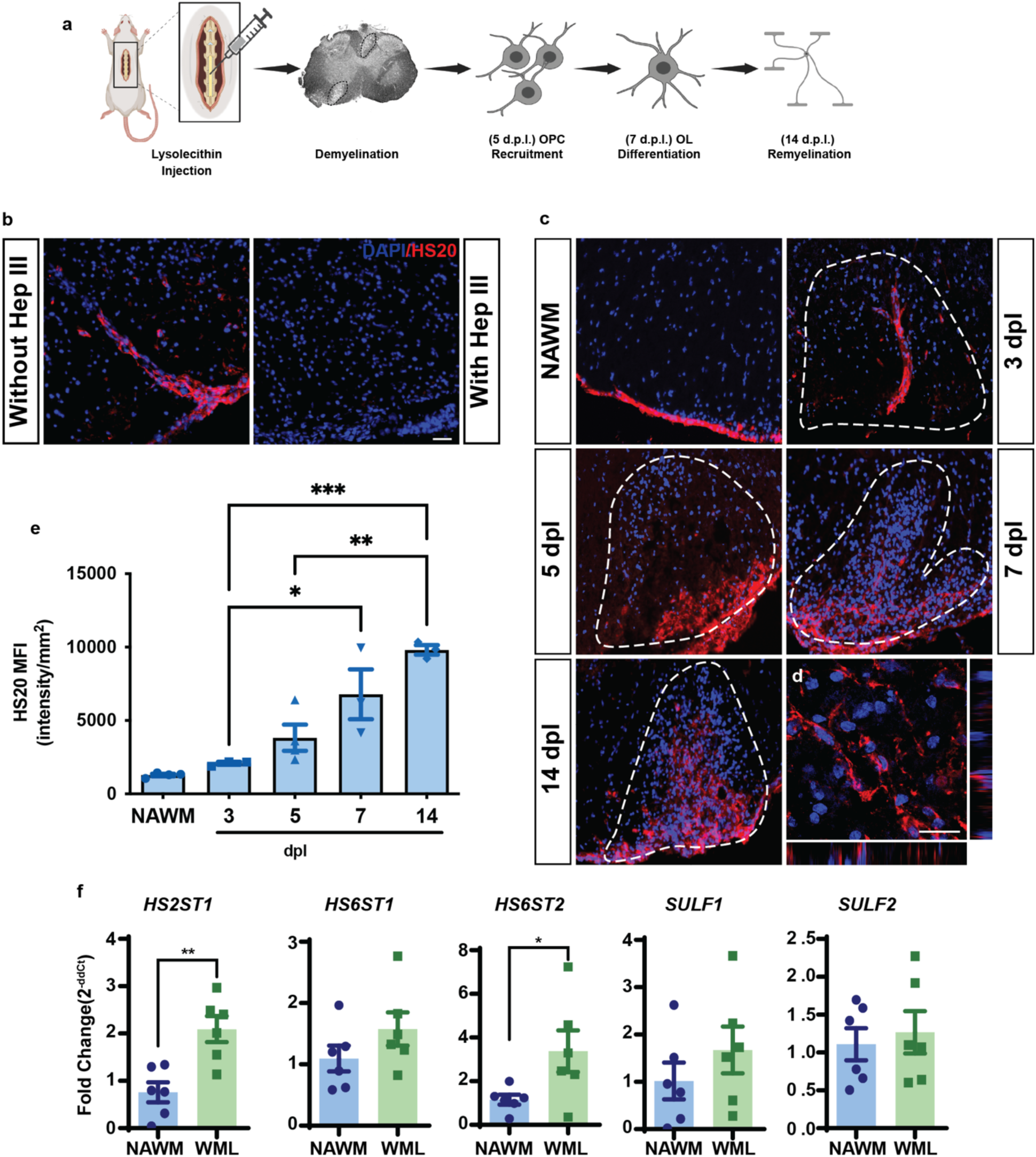
Upregulation of 2-*O* and 6-*O* sulfation following intraspinal lysolecithin-induced demyelination and gene expression of HS-modifying enzymes in human MS lesions. Lysolecithin was injected intraspinally into the ventral white matter of adult mice and heparan sulfate (HS) expression was examined at 3 days post lesion (dpl), 5 dpl, 7 dpl and 14 dpl, that reflects the timeline of oligodendrocyte progenitor cell (OPC) recruitment, differentiation and remyelination, as represented in the schematic (**a**). **b,** 2-*O* and 6-*O* sulfation on the HS were examined using HS20 antibody at 5 dpl. HS20 antibody specificity was validated with Heparin lyase III (Hep III) treatment which eliminated immunoreactivity, co-labeled with DAPI^+^ nuclei (blue). (**c**) HS20 immunofluorescence was assessed in normal-appearing white matter (NAWM), 3 dpl, 5 dpl, 7 dpl and 14 dpl. White dotted lines represent lesion border. HS20 expression was upregulated within the lesion following demyelination. **d**, Orthogonal reconstruction following confocal imaging showed HS20 staining (red) around a subset of DAPI^+^ nuclei (blue) at 14 dpl. **e**, Quantification of HS20 labeling following demyelination (mean ± SEM; *n* = 3-4 mice). HS20 mean fluorescence intensity (MFI) significantly increased from OPC recruitment (3 dpl) through OPC differentiation (5 dpl) and remyelination (14 dpl). One-way ANOVA and Tukey’s post-test, *p < 0.05; **p < 0.01; ***p < 0.001. **f**, Quantitative PCR analysis of *HS2ST1*, *HS6ST1*, *HS6ST2*, *SULF1*, and *SULF2* on mRNA extracted from chronic active MS lesions and NAWM (mean ± SEM, *n* = 6 patients). GAPDH was used as an internal control. Results are expressed as fold change in gene expression in white matter lesions (WML) relative to NAWM controls. Student’s *t*-test: \**p* < 0.05, \*\**p* < 0.01. Scale: 50 µm (**b**); 20 µm (**d**).

To determine whether similar alterations in HS sulfation pathways occur in human disease, we performed qPCR on chronic active multiple sclerosis (MS) lesions and compared them to normal-appearing white matter (NAWM). HS sulfotransferases *HS2ST1* and *HS6ST2* were significantly increased in MS lesions (**Fig. 1f**). *HS2ST1* expression was elevated 2.1 ± 0.3-fold (*p* < 0.01), while *HS6ST2* showed a 3.4 ± 1.0-fold increase (*p* < 0.05; unpaired *t*-tests; *n* = 6 per group). In contrast, expression of *HS6ST1*, *SULF1*, and *SULF2* was not significantly altered (*p* > 0.05). Together, these data demonstrate that HS 2-*O* and 6-*O* sulfation is dynamically upregulated during the differentiation and remyelination phases of toxin-induced demyelination, and the genes potentially responsible for this increase are selectively enriched in human MS lesions. This convergence supports the hypothesis that regulated HS sulfation is conserved across species and may actively contribute to the remyelination signaling milieu.

### Conditional deletion of *Hs2st1* efficiently reduces 2-*O* sulfated HS in OPCs *in vitro* and *in vivo*

To validate conditional deletion of *Hs2st1* in oligodendrocyte progenitor cells (OPCs), primary mouse OPCs were isolated from neonatal mice following tamoxifen-induced recombination and expanded *in vitro* for 7 days. Immunofluorescence analysis revealed a marked reduction in Hs2st1 protein in cKO OPCs compared to wild-type (WT) controls (**Fig. 2a**), while expression of the oligodendrocyte lineage marker Olig2 was preserved. Because *Hs2st1* selectively catalyzes 2-*O* sulfation without affecting N-sulfation, we next assessed whether conditional deletion altered N-sulfated HS. Immunolabeling with the 10E4 antibody showed comparable signal intensity in WT and *Hs2st1* cKO OPCs (**Fig. b**), indicating that loss of *Hs2st1* does not globally perturb N-sulfated HS. Biochemical validation of *Hs2st1* deletion was further confirmed by immunoblotting, which demonstrated a significant reduction in Hs2st1 protein levels in *Hs2st1* cKO OPCs (0.33 ± 0.16-fold relative to WT; *n* = 3; *p* < 0.05, **Fig. 2c,d**).

**Figure 2:**
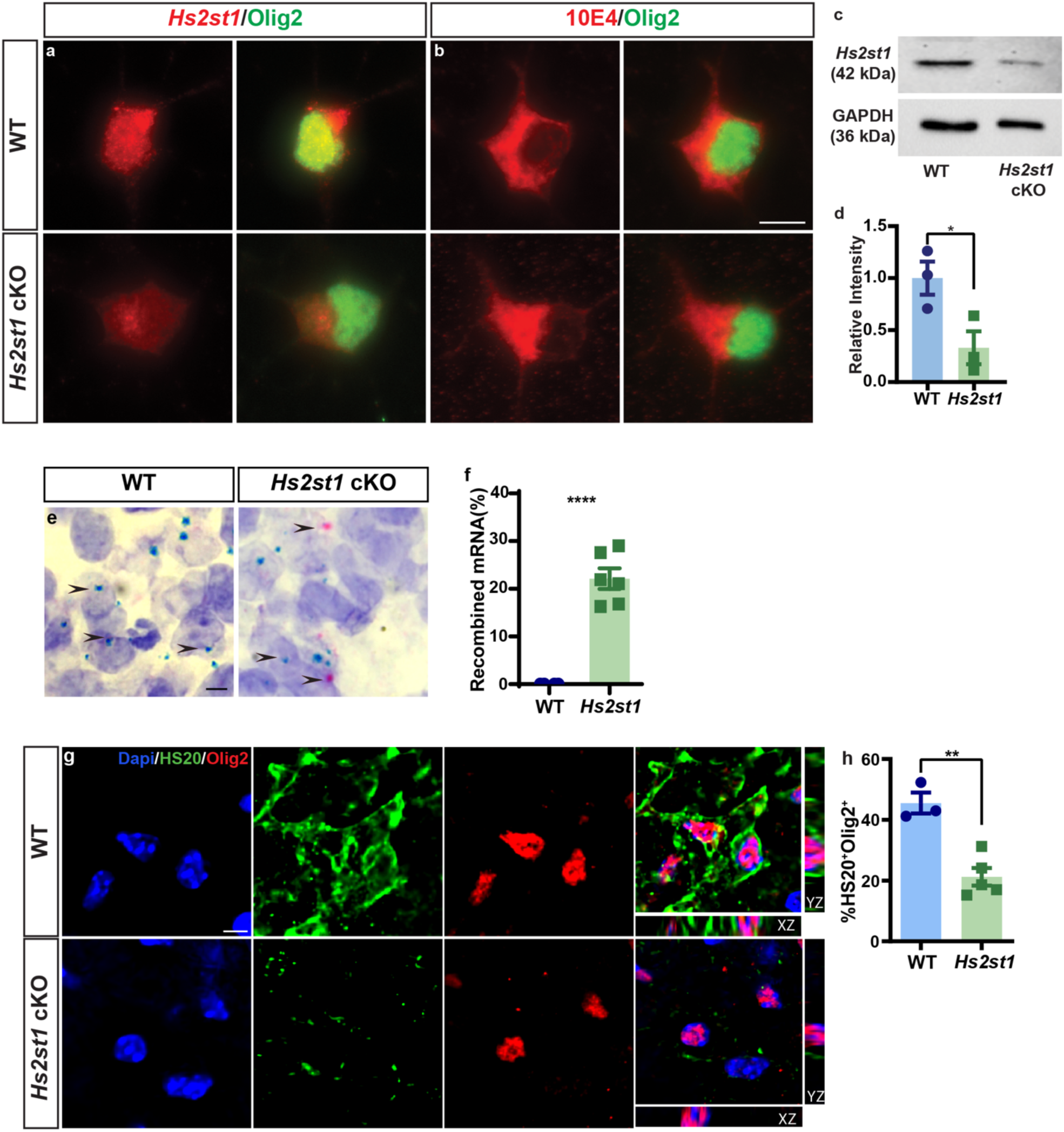
Conditional knockout of OPC-expressed *Hs2st*1 reduces the presence of highly sulfated HSPGs following demyelination. OPC-specific *Hs2st1* conditional knockout was validated in adult *NG2CreERT2: Hs2st1^fl/fl^* mice both *in vitro* and *in vivo*. To validate the conditional knockout *in vitro*, *NG2CreER: Hs2st^fl/^*^fl^ (cre/wt) and corresponding wildtype littermates were injected with 4-hydroxytamoxifen for 4 consecutive days (100 mg/kg IP). mOPCs were isolated using CD140a-based MACS from brains of *Hs2st*1 cKO and WT littermates. After 7 days of cell expansion, knockout was assessed by Hs2st1 (red) and Olig2 (green) immunocytochemistry in WT and *Hs2st1* cKO cells (**a**). Hs2st1 protein (red) is expressed at background levels in mOPCs in the *Hs2st1* cKO. **b**, HS expression was assessed using 10E4 (red) and Olig2 (green) in the WT and *Hs2st1* cKO cells. **c**, Western blot for Hs2st1 in mouse OPCs indicates a significant reduction in Hs2st1 protein expression in cKO cells (**d**) (*n* = 3 samples; * t-test p < 0.05). **e**, Validation of *Hs2st1* cKO using BaseScope probes in adult mice following tamoxifen induction. The distribution of *Hs2st1* mRNA was assessed in WT and *Hs2st1* cKO mice following intraspinal lysolecithin-induced demyelination at 7 dpl. Punctate blue dots and red dots indicate the presence of WT *Hs2st1* and recombined *Hs2st1* mRNA, respectively. **f**, The percentage of cells expressing recombined *Hs2st1* mRNA among the total cells with expressed *Hs2st1* mRNA was determined (*n* = 4 and 6 for WT and *Hs2st1* cKO groups; **** indicates the percentage unpaired t-test *p* = 0.0001). **g**, 2-*O* and 6-*O*-sulfation was identified by HS20 (green) and oligodendroglial lineage cells by Olig2 (red) at 7 dpl. Orthogonal reconstruction showed HS20 expression (green) surrounding Olig2^+^ cells (red). **h,** The percentage of Olig2^+^ cells colocalized with HS20 was quantified and indicated a significant reduction in *Hs2st1* cKO mice (** indicates *t*-test *p* < 0.01). Mean ± SEM shown. Scale: 10 μm (**a**), 50 μm (**e**), and 5 μm (**g**).

To validate recombination *in vivo*, we employed BaseScope *in situ* hybridization using probes specific for WT and recombined *Hs2st1* alleles in spinal cord lesions at 7 dpl. BaseScope signals were detected as discrete puncta within lesion-resident cells (**Fig. 2e**), and probe specificity was confirmed using positive (POLR2A) and negative (dapB) controls (**Supplementary Fig. 2a,b**). Recombined *Hs2st1* probe signal was not detected in WT mice. In contrast, in *Hs2st1* cKO mice, 22.1 ± 2.2% of *Hs2st1*-expressing cells exhibited the recombined allele (*p* < 0.0001; **Fig. 2f**). Given the broad expression of *Hs2st1* mRNA across CNS cell types (Zhang et al., 2014), this mosaic recombination pattern is consistent with selective and efficient OPC-targeted deletion. We next asked whether *Hs2st1* deletion functionally reduced HS 2-*O*/6-*O* sulfation *in vivo*. Immunolabeling with the HS20 antibody revealed a significant reduction in the proportion of HS20⁺ cells among Olig2⁺ oligodendroglial lineage cells within lesions of *Hs2st1* cKO mice compared to WT controls at 7 dpl (**Fig. 2g**). Quantitative analysis showed that only 21.3 ± 2.9% of Olig2⁺ cells were HS20⁺ in *Hs2st1* cKO lesions, compared to 45.5 ± 3.5% in WT mice (*p* < 0.01; **Fig. 2h**). In contrast, the proportion of Olig2⁺ cells positive for N-sulfated HS (10E4) was unchanged between genotypes (**Supplementary Fig. 3**). Together, these findings confirm selective disruption of 2-*O* sulfation without broader alteration of HS sulfation patterns.

### OPC-specific deletion of *Hs2st1* accelerates oligodendrocyte differentiation and remyelination

To determine whether *Hs2st1*-mediated HS sulfation regulates oligodendrocyte lineage dynamics following demyelination, we analyzed lysolecithin-induced spinal cord lesions in OPC-specific *Hs2st1* cKO mice and littermate controls. Lesion centers were identified using solochrome cyanine staining prior to immunofluorescence analysis (**Fig. 3a**). Oligodendroglial lineage cells were quantified using Olig2, and postmitotic mature oligodendrocytes were identified by co-expression of CC1 (**Fig. 3a**). At 7 dpl, *Hs2st1* cKO mice exhibited a marked increase in oligodendrocyte differentiation within lesions. The density of Olig2⁺CC1⁺ postmitotic oligodendrocytes was increased 1.7-fold relative to WT controls (*p* < 0.001; **Fig. 3c**), and the proportion of differentiated oligodendrocytes within the Olig2⁺ lineage population was increased 1.4-fold (*p* < 0.01; **Fig. 3d**; *n* = 5-6 mice per group). This was accompanied by a significant increase in the total density of Olig2⁺ oligodendroglial lineage cells within lesions (*p* < 0.001; **Fig. 3b**), indicating enhanced recruitment and/or expansion of OPCs in the absence of *Hs2st1*.

**Figure 3:**
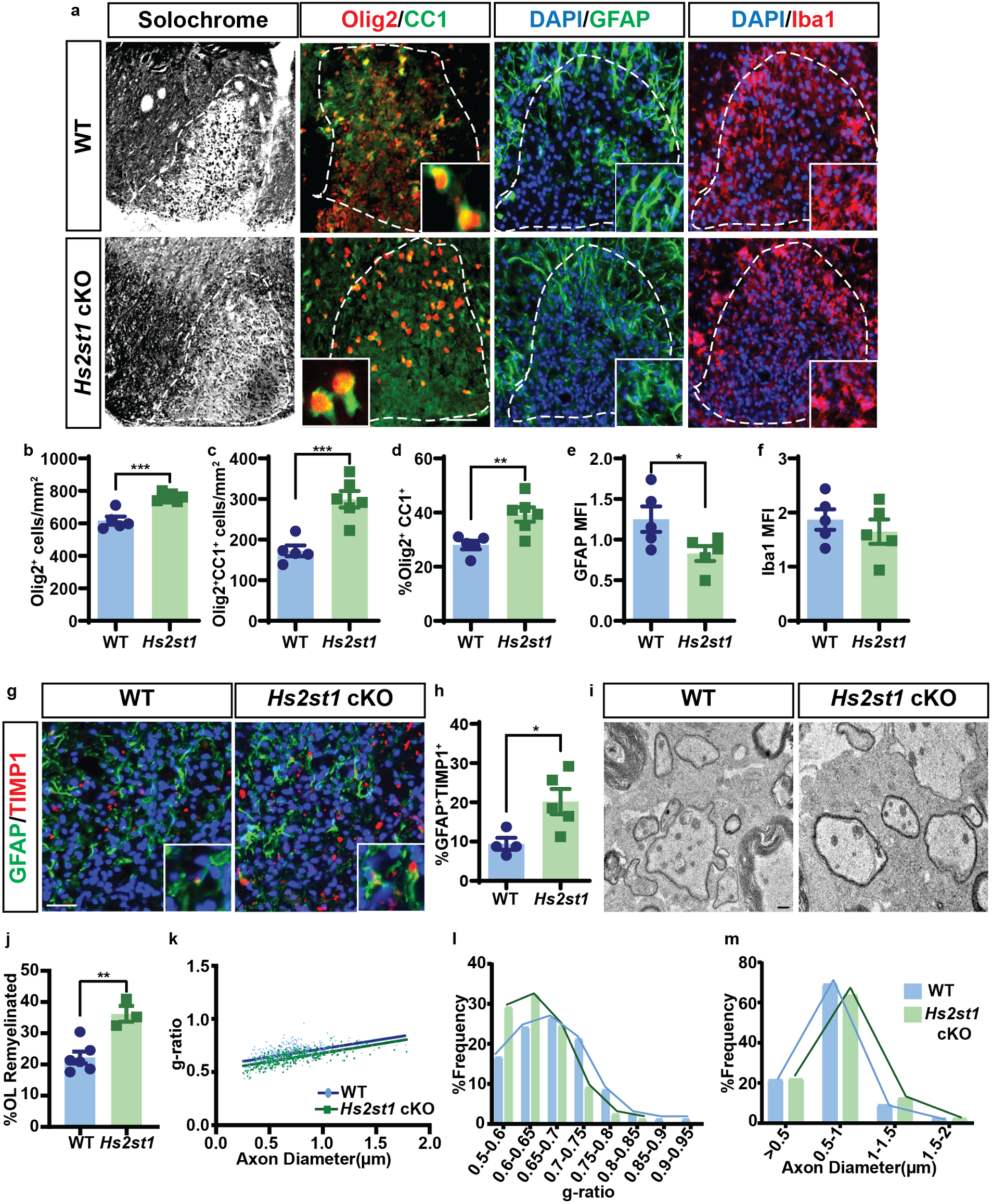
Conditional ablation of *Hs2st*1 in OPCs enhances oligodendrocyte differentiation following focal demyelination. a, Solochrome cyanine staining was performed to visualize demyelinated lesions (7 dpl) in WT and *Hs2st*1 cKO mice. Lesion borders are indicated by a white dotted line. Oligodendroglial lineage cells were identified by Olig2 (red) and mature oligodendrocytes by CC1 (green). Astrogliosis was assessed by GFAP (green), and the microglial response by Iba1 (red). Image inserts show higher magnification to illustrate cell morphology. Quantification of Olig2^+^ (**b**) and CC1^+^ cell density (**c**) (n = 5 and 6 for WT and *Hs2st1*, respectively), and the percentage of CC1^+^ cells within the Olig2^+^ cell population (**d**). *Hs2st*1 cKO significantly enhanced oligodendrocyte density, OPC differentiation and proportion of mature oligodendrocytes. **e**, Mean fluorescence intensity (MFI) of GFAP was significantly reduced, while Iba1 MFI was unaffected in the lesion (**f**) (n = 5 mice). **g**, Astrocytic Timp1 expression was evaluated in WT and *Hs2st1* cKO mice using GFAP and Timp1 markers. **h**, the percentage of GFAP^+^Timp1^+^ cells was quantified among total GFAP^+^ cells at 7 dpl (*n* = 4-5). **i,** Analysis of remyelination by transmission electron microscopy at 14 dpl. **j**, Proportion of remyelinated axons. **k,** Relationship between axon diameter and g-ratio (linear regression shown). Frequency distribution of axonal g-ratio (**l**) and axon diameter (**m**) in lesion (*n* = 6 and 3 mice for WT and *Hs2st1*, respectively, ≥800 axons). Mean ± SEM shown. *, **, *** indicates *t*-test *p* < 0.05, *p* < 0.01, and *p* < 0.001, respectively. Scale: 50 μm (**a, g**) and 1000 nm (**i**).

Because astrocytes and microglia shape the lesion environment during remyelination, we next examined whether OPC-specific *Hs2st1* deletion altered the glial response. At 7 dpl, immunolabeling for GFAP and Iba1 revealed the expected accumulation of reactive astrocytes at lesion borders and amoeboid microglia/macrophages within lesion cores in both genotypes (**Fig. 3a**). Quantification of mean fluorescence intensity (MFI) showed a modest but significant reduction in GFAP signal in *Hs2st1* cKO lesions compared to WT (*p* < 0.05; **Fig. 3e**), whereas Iba1 intensity and distribution were unchanged (**Fig. 3f**). These findings indicate that OPC-specific *Hs2st1* deletion does not grossly alter microglial activation and has only a limited effect on astrocytic reactivity. Given the established role of astrocyte-derived factors in promoting oligodendrocyte differentiation, we next examined expression of the astrocytic secreted protein TIMP-1 (Moore et al., 2011). Co-immunolabeling for GFAP and TIMP-1 revealed a significant increase in the proportion of GFAP⁺TIMP-1⁺ astrocytes in *Hs2st1* cKO lesions relative to WT controls (*p* < 0.05; **Fig. 3g, h**; *n* = 4-5 mice per group), suggesting a permissive trophic environment that may contribute to enhanced oligodendrocyte differentiation.

To assess whether these cellular changes translated into accelerated remyelination, we performed ultrastructural analysis at 14 dpl. *Hs2st1* cKO mice displayed a significantly higher proportion of remyelinated axons within lesions compared to WT controls (*p* < 0.01; **Fig. 3i-j**; *n* = 3-6 mice per group). Consistent with enhanced remyelination, myelin thickness was increased in *Hs2st1* cKO mice, as reflected by a significant reduction in g-ratio (**Fig. 3k**). Linear regression analysis of g-ratio versus axon diameter confirmed a genotype-dependent shift toward thicker myelin across axon calibers. Frequency distribution analysis further demonstrated a reduction in very thinly myelinated axons in *Hs2st1* cKO lesions (two-way ANOVA; genotype *p* < 0.001; **Fig. 3l**). Importantly, axon diameter distributions were unchanged between genotypes (**Fig. 3m**), indicating that *Hs2st1* deletion did not influence axonal swelling. Together, these findings demonstrate that OPC-specific loss of *Hs2st1* accelerates oligodendrocyte recruitment, differentiation, and remyelination following demyelination, identifying HS 2-*O* sulfation as a cell-autonomous negative regulator of myelin repair.

### OPC-specific deletion of *Hs2st1* increases OPC accumulation without altering proliferation or survival

To determine whether the increased oligodendroglial lineage cell density observed in *Hs2st1* cKO mice reflects altered OPC proliferation, we assessed OPC density and cell cycle entry at 5 dpl, a time point at which the majority of Olig2⁺ cells within lysolecithin lesions are OPCs (**Fig. 4**) (Arnett et al., 2004). Mice received EdU injections 24 and 48 h prior to tissue collection to label proliferating cells (**Fig. 4a**). At 5 dpl, the density of Olig2⁺ oligodendrocyte lineage cells within lesions was significantly increased in *Hs2st1* cKO mice compared to WT controls (*p* < 0.01; **Fig. 4b**; *n* = 5 mice per group). However, the density of Olig2⁺EdU^+^ cells incorporating EdU was unchanged between genotypes (**Fig. 4c**; *p* > 0.05), indicating that increased OPC accumulation in *Hs2st1* cKO lesions is not driven by enhanced proliferation. To assess whether differences in cell survival contributed to increased OPC density, we quantified cleaved caspase-3 immunoreactivity within Olig2⁺ cells at 5 dpl (**Fig. 4d**). Cleaved caspase-3 positive debris was comparable between groups (**Fig. 4e**; *n* = 4), indicating that altered apoptosis is unlikely to account for increased OPC accumulation observed in *Hs2st1* cKO lesions.

**Figure 4:**
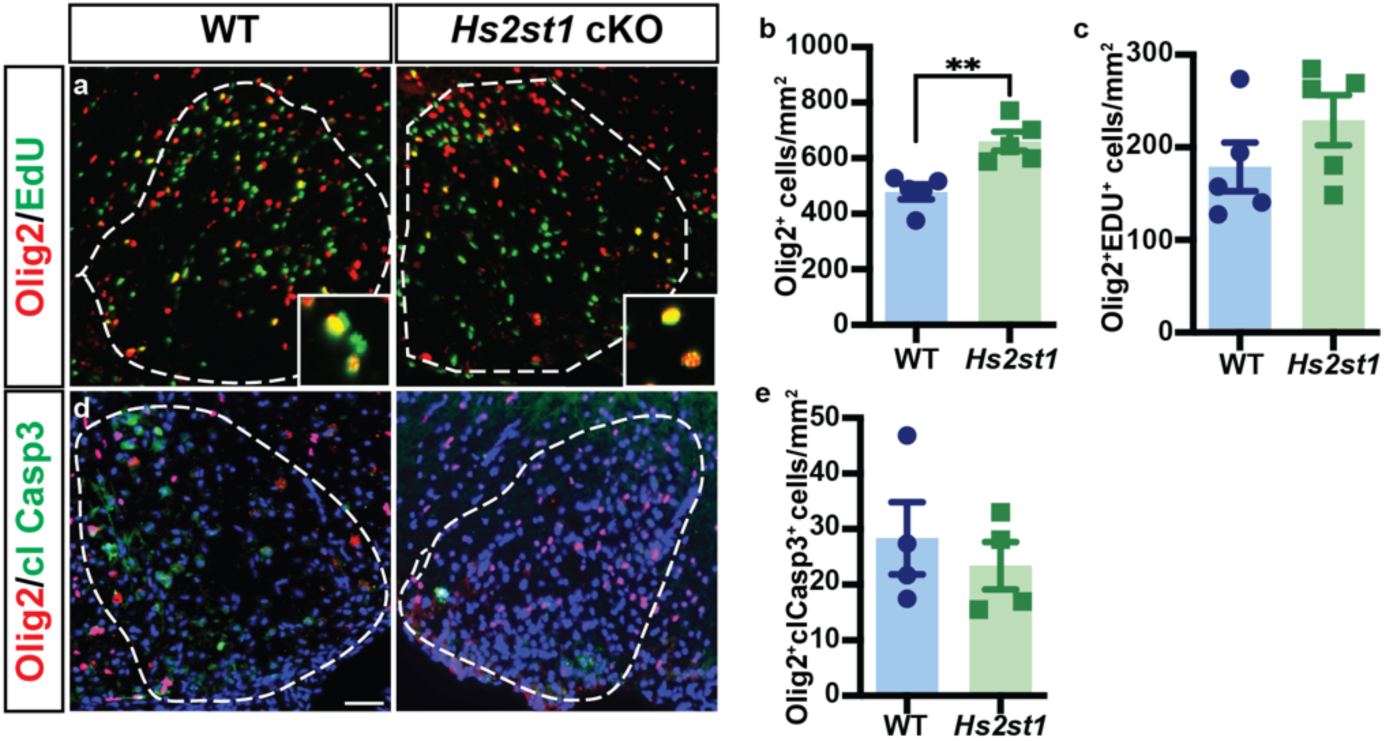
Conditional *Hs2st1* knockout in OPCs accelerates recruitment of oligodendrocyte lineage cells but not proliferation or cell death. The recruitment phase following lysolecithin-induced demyelination was assessed by analyzing WT and *Hs2st1* cKO mice at 5 dpl. To label proliferating cells, EdU was injected 24 and 48 h prior to sacrifice. **a,** Oligodendroglial lineage cells were identified by Olig2 (red) and proliferating cells by EdU (green). Quantification of Olig2^+^ cell density (**b**, *n* = 5 mice/group) and, EdU^+^Olig2^+^ cell density (c, *n* = 4-5 mice). Olig2^+^ cell density was significantly increased in *Hs2st1* cKO mice. **d**, Cell death among oligodendrocyte lineage cells was assessed by colocalization of cleaved caspase 3 (cl casp3; green) with Olig2^+^ (red) cells. **e**, density of Olig2^+^cl Casp3^+^ apoptotic cells (*n* = 4 mice). Mean ± SEM shown. Student’s t-test: \**p* < 0.05. Scale: 50 μm.

Because axonal injury can indirectly influence OPC recruitment and behavior, we next assessed axonal damage using APP immunoreactivity, which accumulates within injured axons in MS (Gehrmann et al., 1995). APP⁺ axonal profiles were readily detected within lesions in both WT and *Hs2st1* cKO mice at 7 dpl (**Supplementary Fig. 4a, c**), but their density did not differ between genotypes (*p* > 0.05; **Supplementary Fig. 4e**). Co-immunolabeling confirmed that APP accumulation was localized to neurofilament-positive axons (**Supplementary Fig. 4b-d**), indicating comparable axonal injury across conditions. Together, these findings demonstrate that OPC-specific deletion of *Hs2st1* increases OPC accumulation within demyelinated lesions without altering OPC proliferation, apoptosis, or axonal injury. These data suggest that reduced HS 2-*O* sulfation primarily enhances OPC recruitment and/or retention within lesions, rather than cell cycle entry or survival.

### Altered HS sulfation suppresses WNT and BMP signaling in OPCs following demyelination

Pathological activation of WNT and BMP signaling pathways inhibits OPC differentiation and limits remyelination efficiency, whereas antagonism of either pathway accelerates oligodendrocyte differentiation and myelin repair (Deininger et al., 1995; Ara et al., 2008; Fancy et al., 2009; Fancy et al., 2011; Sabo et al., 2011). Because 2-*O* and 6-*O* sulfation of heparan sulfate are tightly coupled, such that loss of *Hs2st1* leads to compensatory increases in 6-*O* sulfation (Merry et al., 2001; Merry and Wilson, 2002), we hypothesized that OPC-specific deletion of *Hs2st1* alters responsiveness to 6-*O* sulfation-dependent WNT and BMP signaling that impair differentiation following demyelination.

To test this, we assessed expression of canonical WNT and BMP target genes in OPCs using *in situ* hybridization at 7 dpl. OPCs were identified by *Pdgfra* expression, and target gene expression was quantified within this population (**Fig. 5a**). In *Hs2st1* cKO mice, expression of the WNT target gene *Apcdd1* was significantly reduced, with only 22.2 ± 1.9% of *Pdgfra*⁺ OPCs expressing *Apcdd1*, compared to 39.0 ± 1.3% in WT controls (*p* < 0.0001; **Fig. 5b**; *n* = 5-6 mice per group). Similarly, expression of the BMP target gene *Id4* was significantly decreased in *Hs2st1* cKO OPCs (15.5 ± 1.7%) relative to WT (27.0 ± 2.5%; *p* < 0.01; **Fig. 5c**; *n* = 4-5 mice per group). These data indicate that OPC-specific loss of *Hs2st1* attenuates both WNT and BMP signaling within OPCs during remyelination, consistent with a pro-differentiative shift in the lesion environment.

**Figure 5:**
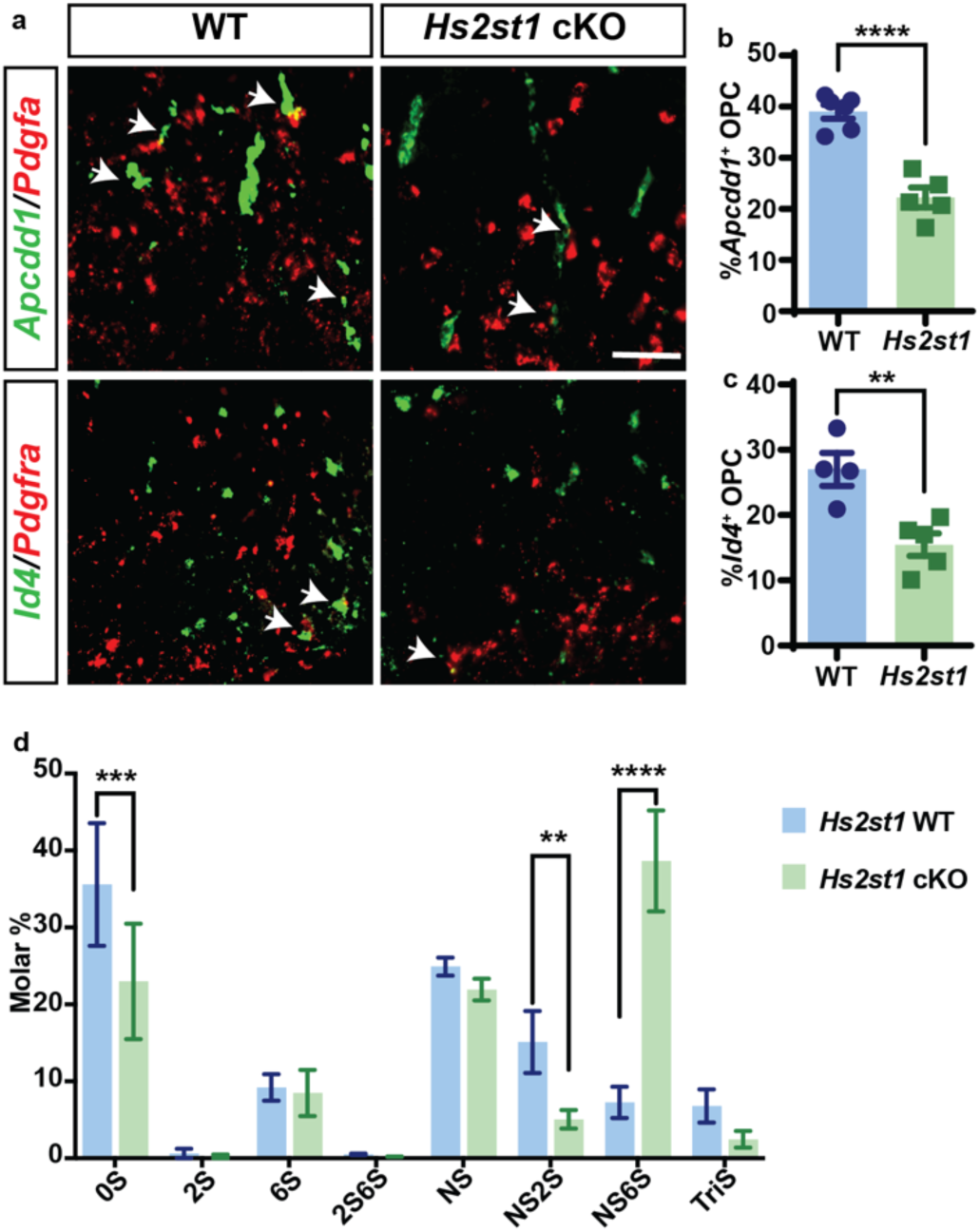
*Hs2st1* cKO accelerated OPC differentiation by inhibition of the WNT/BMP signaling pathway and increasing levels of highly sulfated HS. a, RNAscope *in situ* hybridization images demonstrates co-localization of OPC-expressed *Pdgfra* (red) with WNT target gene *Apcdd1* (green) and BMP target gene *Id4* (green) at 7 dpl. Arrows indicate double-labeled OPCs. The percentage of *Apcdd1* (**b**) and *Id4* (**c**) co-labeled *Pdgfra*^+^ OPCs was determined (*n* = 4-6 mice). **d**, Relative abundance of cell surface HS disaccharides from OPCs isolated from WT and *Hs2st1* cKO mice was analyzed using liquid chromatography-tandem mass spectrometry (*n* = 3 independent biological samples). Mean ± SEM shown. Significance in **b** and **c** was determined using *t*-tests and in **d** using Sidak’s multiple comparison test. **, ***, and **** indicate *p* < 0.01, *p* < 0.001, and *p* < 0.0001, respectively. Scale: 50 µm.

### Loss of *Hs2st1* induces compensatory enrichment of 6-*O* sulfated HS species in OPCs

To determine whether altered WNT and BMP signaling in *Hs2st1* cKO OPCs is associated with changes in HS sulfation composition, we performed quantitative disaccharide profiling of cell-surface HS isolated from OPCs using liquid chromatography-tandem mass spectrometry (LC-MS/MS). HS disaccharides were resolved using eight defined standards, including unsulfated (0S), mono-sulfated (2S, 6S, NS), di-sulfated (NS2S, NS6S, 2S6S), and tri-sulfated (TriS) species (**Fig. 5d**).

As expected, disaccharide compositional analysis demonstrated a reduction in 2-*O*-sulfated species in *Hs2st1* cKO OPCs. Notably, this was accompanied by a striking 5.3-fold increase in the abundance of ΔUA-GlcNS6S (NS6S), a disaccharide containing both N-sulfation and 6-*O* sulfation, in *Hs2st1* cKO OPCs (38.6 ± 6.6%) compared to WT controls (7.2 ± 2.0%, *p* < 0.0001). This shift is consistent with compensatory enhancement of 6-*O* sulfation in the absence of 2-*O* sulfation. Statistical analysis across disaccharide classes confirmed a significant genotype-dependent alteration in HS sulfation composition (two-way ANOVA; *n* = 3 independent biological replicates). Together, these data demonstrate that *Hs2st1* deletion drives a selective remodeling of HS sulfation toward 6-*O* enriched HSPG in OPCs, providing a biochemical mechanism by which altered HS composition may dampen WNT and BMP signaling and promote oligodendrocyte differentiation.

### Loss of 6-*O* sulfotransferase activity restricts OPC recruitment and proliferation following demyelination

Our previous studies demonstrated that removal of 6-*O* sulfation by sulfatases generates an inhibitory signaling environment following demyelination (Saraswat et al., 2021b). Because loss of *Hs2st1* leads to compensatory increases in OPC 6-*O* sulfation, we hypothesized that reducing 6-*O* sulfation through ablation of 6-*O* sulfotransferases, either alone or in combination with loss of 2-*O* sulfation, would exacerbate inhibitory signaling and impair OPC recruitment following demyelination. To test this, we analyzed lysolecithin-induced spinal cord lesions at 5 dpl in four NG2CreER-based mouse models targeting HS *O*-sulfation: *Hs6st1^fl/fl^*, *Hs6st2^−/−^*, *Hs6st1^fl/fl^;Hs6st2^−/−^*, and *Hs2st1^fl/fl^;Hs6st1^fl/fl^;Hs6st2^−/−^* (**Fig. 6**). Because a conditional *Hs6st2* allele is not currently available and single-cell transcriptomic datasets demonstrate selective enrichment of *Hs6st2* expression in OPCs (Zheng et al., 2025), we used a germline *Hs6st2* null allele as a proxy for OPC-targeted loss of 6-*O* sulfotransferase activity. Oligodendrocyte lineage cells were quantified using Olig2, and newly generated OPCs were assessed as the density of EdU⁺Olig2⁺ cells within lesions.

**Figure 6:**
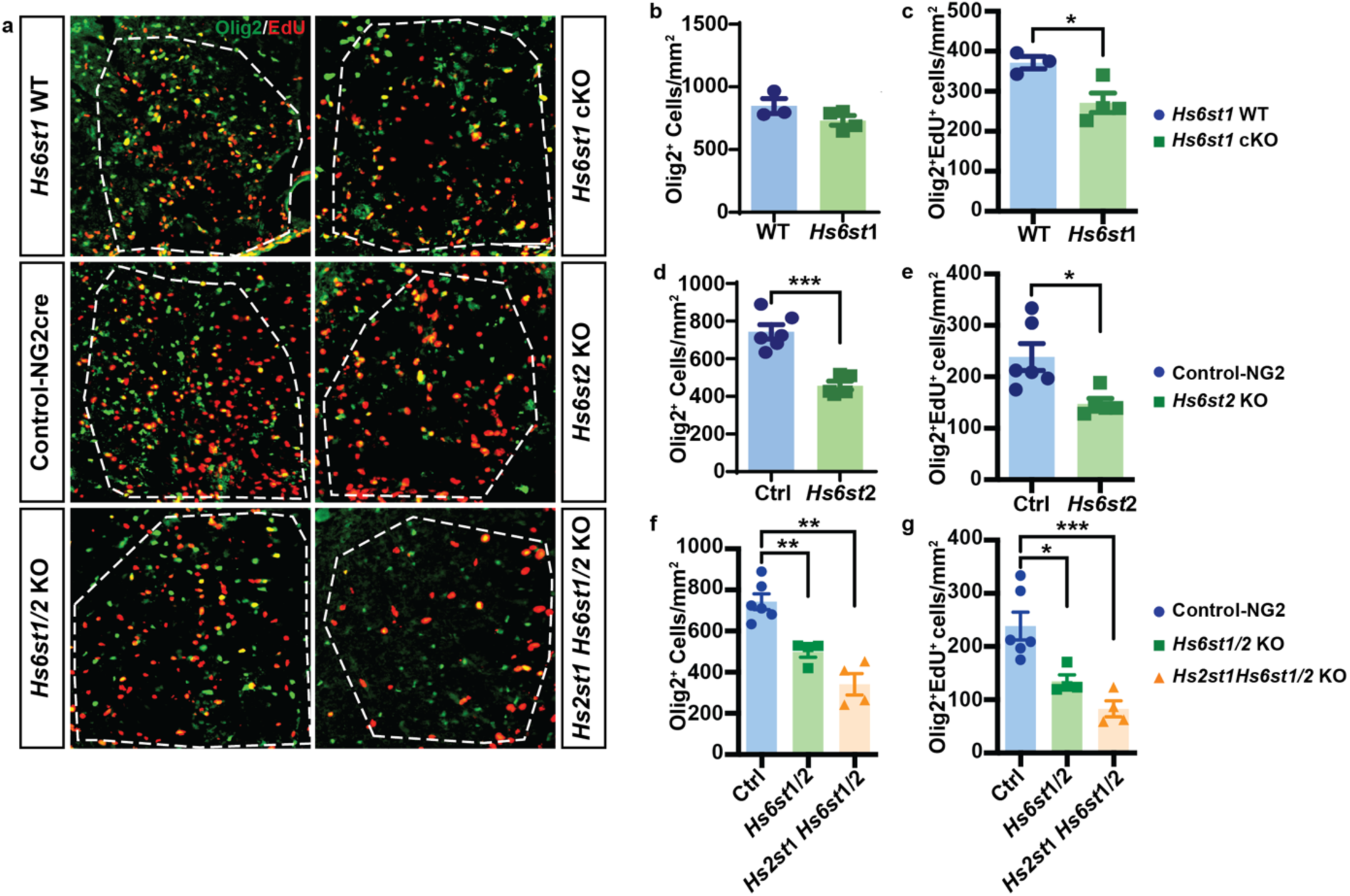
Reduced 6-*O* sulfation by *Hs6st1/2* KO reduced OPC recruitment and proliferation which could not be rescued by concurrent *Hs2st1* deletion following demyelination. To determine the contribution of 6-*O* sulfation to OPC recruitment and proliferation, we induced OPC-specific deletion of *Hs6st1*, germline deletion of *Hs6st2,* or the combination of both. Additionally, the potential interaction with 2-*O* sulfation was determined by combined OPC-specific deletion of *Hs2st1* in 6-*O* deficient *Hs6st1/2* mutant mice. **a**, OPC recruitment was assessed at 5 dpl in the spinal cord lysolecithin model. Oligodendroglial lineage cells were identified by Olig2 (red) and proliferating cells by EdU^+^ (green). The density of Olig2^+^ cells and proliferating Olig2^+^EdU^+^ OPCs in *Hs6st1* cKO (**b, c**, *n* = 3 and 4, WT and cKO, respectively), and *Hs6st2* KO was quantified (**d**, **e**, *n* = 6 and 5 for WT and KO, respectively). *Hs6st1* cKO did not alter the overall OPC density but significantly reduced the number of proliferating OPCs, whereas both the density of total and proliferating Olig2^+^ OPCs was significantly decreased in the *Hs6st2* KO mice (*n* = 6 and 5 for control and *Hs6st2* KO, respectively). Double knockout of both 6-*O* sulfotransferases (*Hs6st1^fl/fl^; Hs6st2^−/−^*) similarly resulted in reduction of Olig2^+^ cell density (**f**) and proliferating OPCs (**g**). The combined deletion of *Hs2st1* in the context of 6-*O* depletion did not rescue the observed reduced OPC density (**f, g**, *n* = 6, 4, and 4 for WT, *Hs6st1/2*, and *Hs2st1; Hs6st1/2*, respectively). One-way ANOVA; *, **, and *** indicate Dunnett’s post-test *p* < 0.05, *p* < 0.01, and *p* < 0.001. Mean ± SEM shown. Scale: 50 µm.

Conditional deletion of *Hs6st1* alone did not significantly alter the density of Olig2⁺ oligodendroglial lineage cells within lesions (*p* > 0.05; **Fig. 6b**; *n* = 3-4 mice per group) but resulted in a significant reduction in the density of proliferating OPCs (*p* < 0.05; **Fig. 6c**), indicating a contribution of *Hs6st1* to OPC cell-cycle entry. In contrast, loss of *Hs6st2* caused a pronounced reduction in both total Olig2⁺ cell density (*p* < 0.001; **Fig. 6d**) and EdU^+^Olig2^+^ OPC density (*p* < 0.05; **Fig. 6e**), demonstrating a critical role for *Hs6st2* in supporting OPC recruitment and proliferation following demyelination. Consistent with these findings, combined deletion of *Hs6st1* and *Hs6st2* resulted in marked reductions in Olig2⁺ cell density (one-way ANOVA; F(2, 8.2) = 27.3; **Fig. 6f**) and EdU⁺Olig2⁺ OPC density (**Fig. 6g**). Importantly, concurrent deletion of *Hs2st1* in the *Hs6st1/2*-deficient background failed to rescue deficits in either OPC accumulation or proliferation, indicating that loss of 2-*O* sulfation does not compensate for impaired OPC responses caused by reduced 6-*O* sulfation. Together, these data demonstrate that 6-*O* sulfation of heparan sulfate proteoglycans is required to support both OPC recruitment and proliferative expansion following demyelination, and that this function cannot be restored by additional loss of 2-*O* sulfation.

### 6-*O* sulfation is required for efficient oligodendrocyte differentiation following demyelination

To determine whether disruption of HS *O*-sulfation alters oligodendrocyte differentiation during remyelination, we quantified oligodendrocyte lineage cells (Olig2⁺) and mature oligodendrocytes (Olig2⁺CC1⁺) within lysolecithin-induced spinal cord lesions at 7 dpl in HS 2-*O* and 6-*O* deficient mutant mice (**Fig. 7a**). Deletion of *Hs6st1* alone did not significantly affect the density of Olig2⁺ lineage cells or Olig2⁺CC1⁺ mature oligodendrocytes compared to controls (*p* = 0.5 for both; **Fig. 7b,c**; *n* = 5-9 mice per group). Likewise, the proportion of mature oligodendrocytes among the Olig2⁺ population was unchanged (*p* = 0.57), indicating that *Hs6st1* is dispensable for oligodendrocyte differentiation in this context.

**Figure 7:**
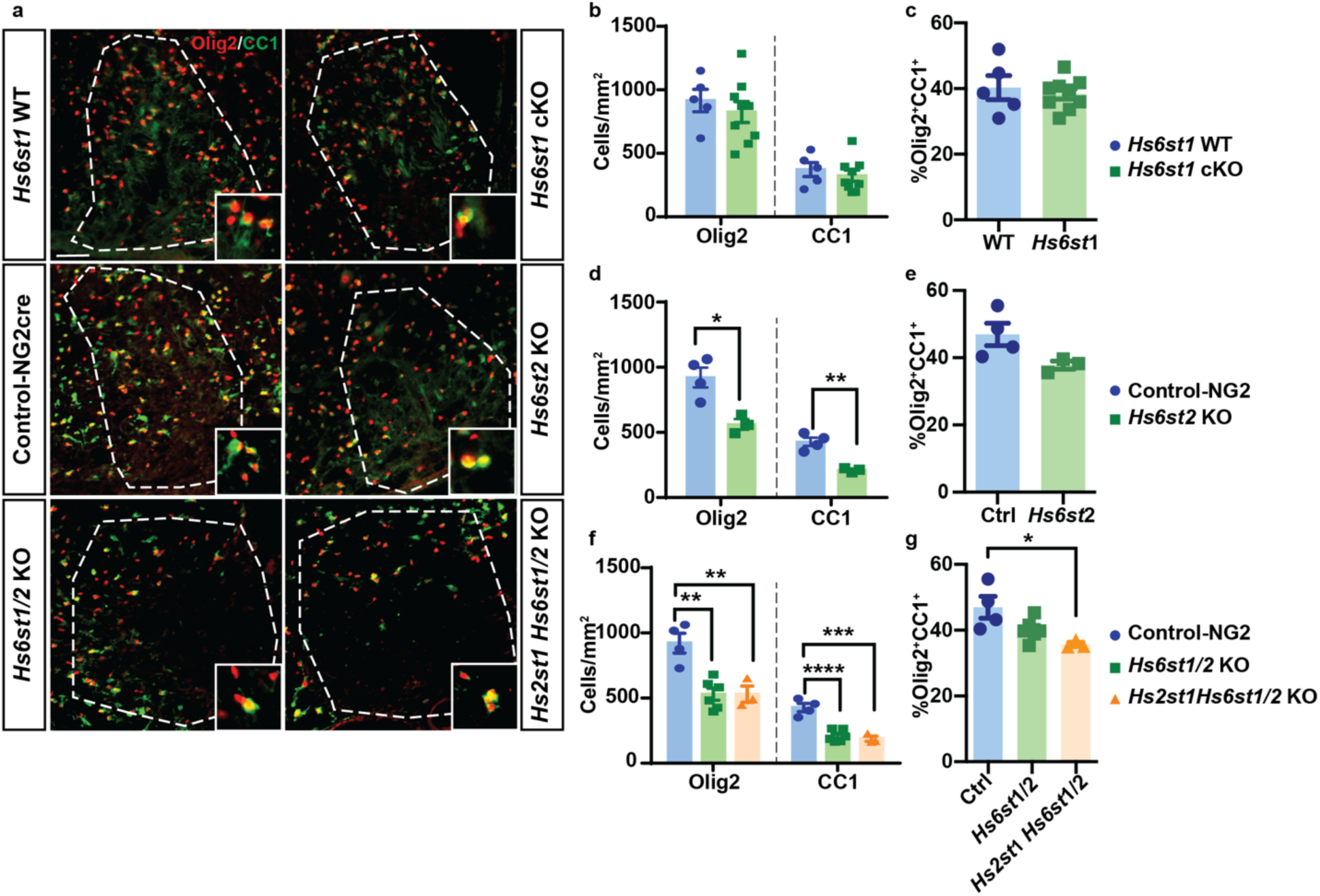
Reduced 6-*O* sulfation inhibits OPC differentiation following focal demyelination independent of 2-*O* sulfation. The role of 6-*O* sulfation in OPC differentiation following demyelination was investigated using lysolecithin-induced demyelination at 7 dpl. **a**, Oligodendroglial lineage cells were identified by Olig2 (red) and mature oligodendrocytes by CC1 (green). The Olig2^+^ and CC1^+^ cell density (**b, d, f**), and the percentage of CC1^+^ cells within the Olig2^+^ cell population (**c, e, g**) was quantified in *Hs6st1* cKO, *Hs6st2* KO, 6-*O* deficient *Hs6st1/2* double mutant, and 6/2-*O* deficient *Hs2st1; Hs6st1/2* triple mutant mice and corresponding controls. *Hs6st1* cKO had no detectable effect on oligodendroglial density or differentiation at 7 dpl. In contrast, *Hs6st2* KO and *Hs6st1/2* double knockout results in decreased Olig2^+^ and CC1^+^ cell density. Importantly, *Hs2st1* cKO in the 6-*O* deficient *Hs6st1/2* did not rescue the reduced density of Olig2^+^ oligodendrocyte lineage and CC1^+^Olig2^+^ oligodendrocytes. One-way ANOVA; *, **, *** and **** indicate Dunnett’s post-test *p* < 0.05, *p* < 0.01, *p* < 0.001, and *p* < 0.0001, respectively. Mean ± SEM shown. Scale: 50 µm.

In contrast, deletion of *Hs6st2* significantly reduced both total Olig2⁺ lineage cell density (*p* < 0.05; **Fig. 7d**) and Olig2⁺CC1⁺ mature oligodendrocyte density (*p* < 0.01), while the proportion of differentiated cells among the remaining Olig2⁺ population was not significantly altered (**Fig. 7e**). Importantly, analysis of adult spinal cord white matter prior to demyelination revealed no differences in oligodendrocyte lineage density or differentiation in *Hs6st2*-null mice compared to controls (**Supplementary Fig. 5**), indicating that these deficits reflect impaired regenerative responses rather than a developmental phenotype. Combined deletion of *Hs6st1* and *Hs6st2* recapitulated the reduction in Olig2⁺ cells and mature oligodendrocytes observed in *Hs6st2* mutants (one-way ANOVA; **Fig. 7f**). Together, these findings indicate that loss of 6-*O* sulfotransferase activity, driven predominantly by *Hs6st2* deletion, impairs oligodendrocyte accumulation and differentiation during the regenerative phase of demyelination.

Importantly, concurrent deletion of *Hs2st1* in the *Hs6st1/2*-deficient background failed to rescue these deficits and, in some measures, further reduced the proportion of mature oligodendrocytes (**Fig. 7g**). Thus, loss of 2-*O* sulfation does not compensate for impaired 6-*O* sulfation during oligodendrocyte differentiation.

### Loss of HS 2-*O* and 6-*O* sulfation does not alter reactive gliosis following demyelination

Reactive astrocytes and microglia contribute to the inflammatory milieu that shapes remyelination efficiency. To determine whether the OPC phenotypes observed in HS sulfotransferase mutants reflect altered gliotic responses, we examined astrocyte and microglial activation at 7 dpl in HS 2-*O* and 6-*O*-deficient mutant mice. Immunofluorescence for GFAP revealed robust astrocyte activation at lesion borders across all genotypes (**Fig. 8a**). Quantification of GFAP MFI showed no significant differences between groups (one-way ANOVA; *F*(3,18) = 1.948; **Fig. 8b**; *n* = 4-8 mice per group). Similarly, Iba1⁺ microglia/ macrophages were distributed throughout lesion cores and borders in all genotypes (**Fig. 8c**), and Iba1 MFI did not differ significantly between groups (one-way ANOVA; *F*(3,18) = 1.011; **Fig. 8d**). These findings indicate that genetic disruption of 2-*O* and/or 6-*O* sulfation in OPCs does not grossly alter astrocytic or microglial responses following demyelination, supporting a predominantly OPC-intrinsic mechanism underlying the observed effects on remyelination.

**Figure 8:**
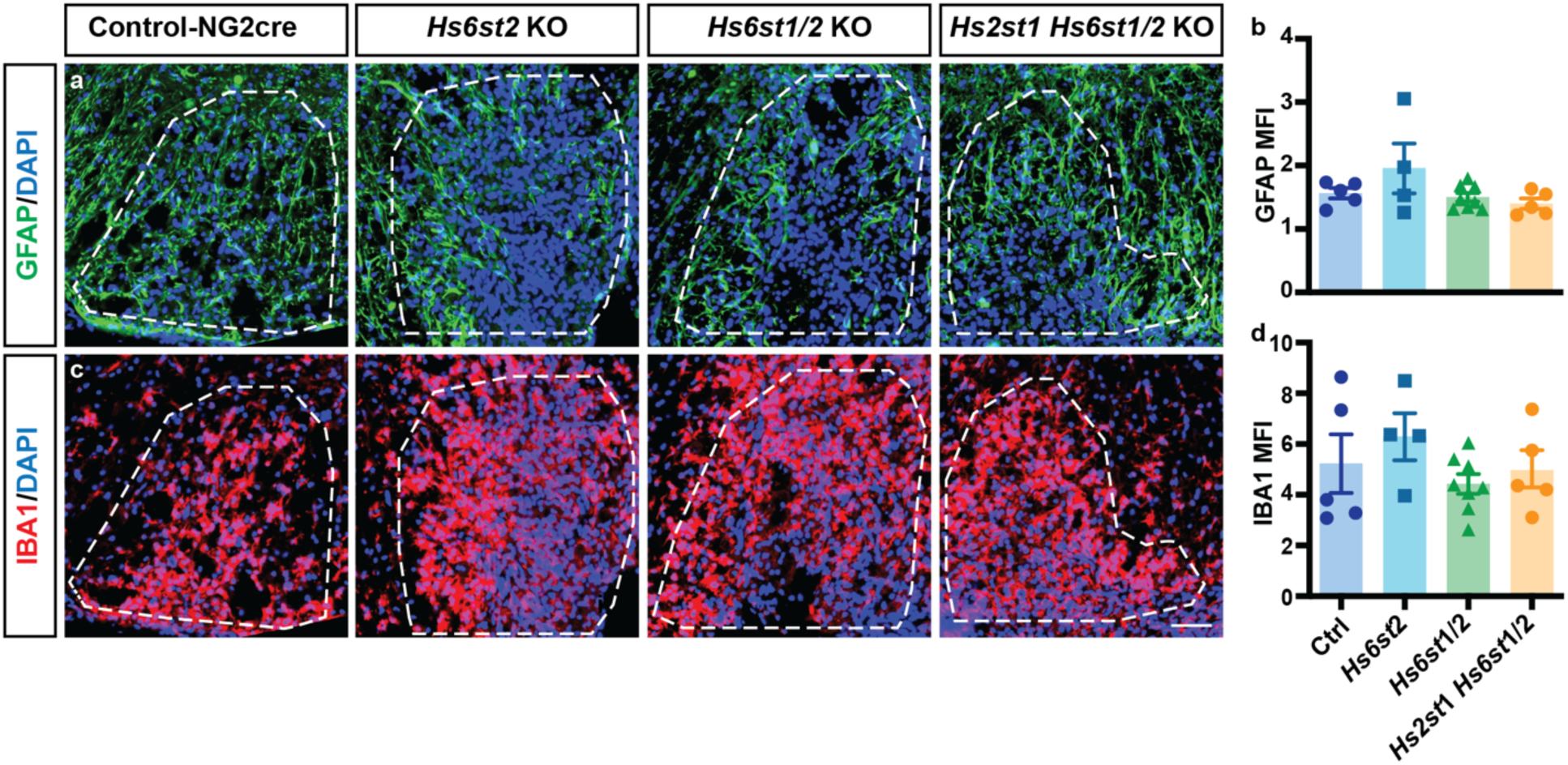
Heparan sulfate 6-*O* sulfation does not alter the innate inflammatory response of astrocytes and microglia following focal demyelination. The innate immune response of astrocytes and microglia was evaluated in control, *Hs6st2^−/−^*, *Hs6st1/2* and *Hs2st1; Hs6st1/2* mice at 7 dpl. **a**, Astrocyte reactivity was assessed using GFAP (green) and mean fluorescence intensity (MFI) was quantified within the lesion region (**b**). The microglial response was examined using Iba1 (red, **c**) and MFI was quantified (**d**). There were no significant changes across groups (*n* = 5, 4, 8, and 5 mice for control, *Hs6st2^−/−^*, *Hs6st1/2* (cKO/KO), and *Hs2st1; Hs6st1/2* (cKO/KO), respectively; one-way ANOVA). Mean ± SEM shown. Scale: 50 µm.

### Reduction of 6-*O* sulfotransferase activity does not suppress WNT or BMP signaling in OPCs

Extracellular sulfatases promote inhibitory WNT and BMP signaling in OPCs following demyelination, and their deletion enhances remyelination by attenuating pathway activation. Because 6-*O* sulfation status influences ligand-receptor interactions within these pathways, we asked whether genetic reduction of 6-*O* sulfation through deletion of *Hs6st* enzymes alters WNT or BMP signaling activity in OPCs during remyelination. To test this, we quantified expression of the canonical WNT target gene *Apcdd1* and the BMP target gene *Id4* in *Pdgfra*⁺ OPCs using multicolor fluorescence *in situ* hybridization at 7 dpl in HS 6-*O* deficient mutant mice (**Fig. 9**). In contrast to the suppression of WNT and BMP targets observed following *Hs2st1* deletion, no significant differences were detected in the proportion of *Apcdd1*⁺ OPCs among 6-*O* deficient genotypes compared to controls (one-way ANOVA; *F*(3,10) = 0.9308; **Fig. 9a,b**; *n* = 3-4 mice per group). Similarly, *Id4* expression within *Pdgfra*⁺ OPCs was unchanged across genotypes (one-way ANOVA; *F*(3,12) = 2.177; **Fig. 9c-d**; *n* = 3-5 mice per group). These findings indicate that, unlike loss of *Hs2st1*, reduction of 6-*O* sulfotransferase activity does not alter canonical WNT or BMP signaling in OPCs at 7 dpl. Thus, the impaired recruitment and differentiation observed in 6-*O* deficient mutants are unlikely to be mediated through altered WNT/BMP transcriptional activity at this stage of remyelination.

**Figure 9:**
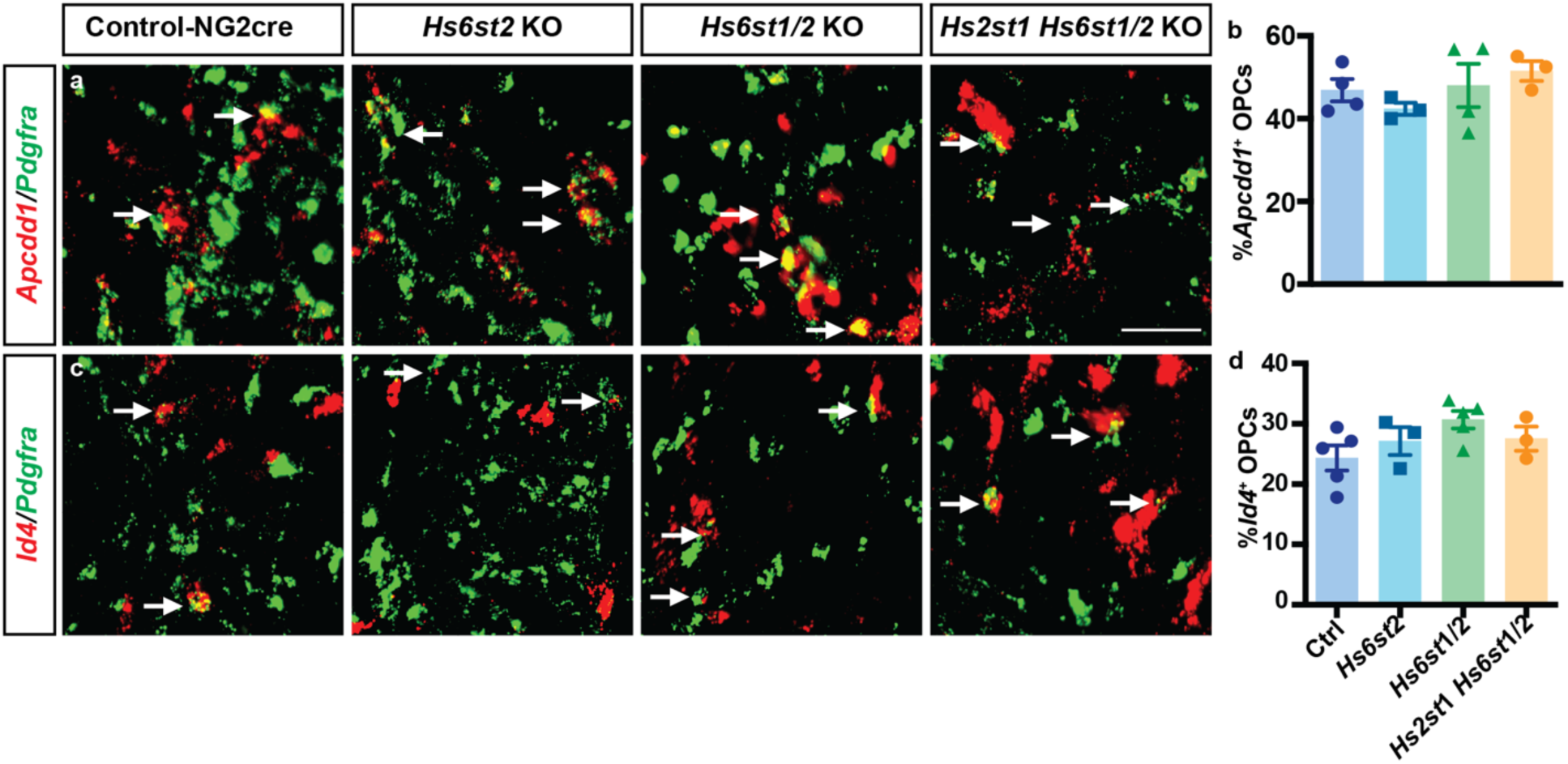
6-*O* sulfotransferase expression is required for the attenuation of WNT and BMP signaling in OPCs observed following *Hs2st1* cKO. RNAscope *in situ* hybridization was used to evaluate the WNT/BMP signaling pathway in control, 6-*O*-deficient *Hs6st2* and *Hs6st1/2* mice, and combined 6-*O-* and 2-*O*-deficient *Hs2st1;Hs6st1/2* mice following lysolecithin-induced demyelination at 7 dpl. Arrows indicate co-localization of *Pdgfra*^+^ OPCs (green) with WNT target gene *Apcdd1* (red, **a**) and BMP target gene *Id4* (red, **c**). Quantification of WNT pathway activity (*Apcdd1*%, **b**) and BMP pathway activity (*Id4*%, **d**) among *Pdgfra*^+^ OPCs was performed. Mean ± SEM is shown. Scale: 50 µm.

## DISCUSSION

Failure of remyelination in multiple sclerosis (MS) is driven not only by intrinsic deficits in oligodendrocyte progenitor cells (OPCs) (Franklin et al., 2024), but also by persistent alterations in the extracellular matrix (ECM) that reshape the lesion microenvironment (Lau et al., 2013; Ghorbani and Yong, 2021). Among these ECM components, heparan sulfate (HS) proteoglycans undergo dynamic changes in their synthesis and sulfation pattern during injury and repair (Basu et al., 2022), and serve as critical determinants of growth factor signaling (Sarrazin et al., 2011). However, how specific *O*-sulfation patterns regulate OPC behavior during remyelination has remained unclear. Here, we demonstrate that HS *O*-sulfation constitutes a hierarchical regulator of myelin repair in which 6-*O* sulfation plays a dominant, permissive role in supporting OPC recruitment and differentiation, while 2-*O* sulfation modulates this process indirectly through its influence on 6-*O* sulfation levels. Genetic ablation of *Hs2st1* in OPCs enhances remyelination, suppresses canonical WNT and BMP signaling, and increases 6-*O* sulfated disaccharides, phenocopying the pro-repair effects observed following sulfatase inhibition. In contrast, deletion of *Hs6st2* impairs oligodendrocyte accumulation and differentiation without altering WNT or BMP target expression, and combined loss of 6-*O* sulfation abolishes the pro-remyelinating effects of *Hs2st1* deletion. Together, these findings identify 6-*O* sulfation as a key extracellular determinant of OPC regenerative competence and position 2-*O* sulfation as a modulatory node that shapes remyelination through secondary regulation of 6-*O* sulfation.

Our findings demonstrate that HS sulfation is not static within demyelinating lesions but is dynamically regulated across the phases of repair, consistent with the broader concept that proteoglycan sulfation is altered during injury and aging (Fawcett and Kwok, 2022). In the lysolecithin model, highly sulfated HS epitopes increased during the differentiation and remyelination stages, coinciding temporally with OPC maturation and myelin formation. Consistent with this, transcripts encoding HS-modifying enzymes, including *HS2ST1* and *HS6ST2*, were enriched in chronic active MS lesions relative to normal-appearing white matter, suggesting that analogous sulfation remodeling occurs in human disease. Published single-cell and single-nucleus transcriptomic datasets further support the translational relevance of these findings. Across independent analyses of human MS tissue, *HS2ST1* expression was detected in OPCs and early differentiating oligodendrocytes but was also present in astrocytes and other glial populations, whereas *HS6ST2* expression was largely restricted to OPCs and committed oligodendrocyte precursors (Jakel et al., 2019; Trobisch et al., 2022). Similar lineage-restricted expression of *Hs6st2* was observed in mouse demyelination datasets, while *Hs2st1* exhibited broader cellular expression (Pandey et al., 2022; Zheng et al., 2025). Furthermore, transcriptomic analysis of purified human OPCs confirmed expression of *HS2ST1*, *HS6ST1*, and *HS6ST2*, supporting conservation of the enzymatic machinery responsible for HS 2-*O* and 6-*O* sulfation within the human oligodendrocyte lineage. Together, these observations suggest that lesion-associated ECM remodeling includes active regulation of HS *O*-sulfation rather than simple accumulation of proteoglycans and suggest that regulated sulfation patterns are conserved across species. Earlier work in rat OPCs demonstrated that glial activation following injury is accompanied by increased 2-*O* sulfation and elevated *Hs2st1* expression (Properzi et al., 2008), supporting the concept that HS sulfation is actively regulated within the injured CNS. This contrasts with the accumulation of basement membrane-associated HSPGs observed within inflammatory cuffs in MS, which more likely influence immune cell infiltration and inflammatory signaling (van Horssen et al., 2005; van Horssen et al., 2006). The temporal alignment between increased sulfation and OPC differentiation further implies that specific sulfation motifs contribute functionally to the regenerative microenvironment and provides a rationale for dissecting the individual roles of 2-*O* and 6-*O* sulfation in remyelination.

A central finding of this study is that 6-*O* sulfation represents the dominant HS modification supporting oligodendrocyte recruitment and differentiation during remyelination. Genetic ablation of *Hs6st2* significantly reduced the accumulation of Olig2⁺ lineage cells and mature oligodendrocytes within lesions. In contrast, deletion of *Hs6st1* alone did not alter overall oligodendrocyte lineage density or subsequent differentiation, although a transient reduction in the density of proliferating OPCs was observed at 5 days post lesion. Combined loss of *Hs6st1* and *Hs6st2* did not exacerbate the *Hs6st2* phenotype. Moreover, concurrent deletion of *Hs2st1* failed to rescue the deficits observed in 6-*O*-deficient mice, indicating that 2-*O* sulfation cannot compensate for impaired 6-*O* sulfation during repair. Together, these data identify *Hs6st2* as the primary 6-*O* sulfotransferase governing OPC regenerative competence in the adult CNS, with *Hs6st1* contributing only modestly to alter early progenitor dynamics. Consistent with this genetic hierarchy, re-analysis of publicly available single-cell RNA-sequencing datasets revealed selective enrichment of *Hs6st2* transcripts in OPCs in the uninjured mouse brain and at 5 days following injection of lysolecithin (Pandey et al., 2022), whereas both *Hs2st1* and *Hs6st1* were broadly expressed across neural cell types. Similarly, human MS datasets demonstrate enrichment of *HS6ST2* within OPC clusters (Jakel et al., 2019; Trobisch et al., 2022). Together, these genetic and transcriptional findings support a hierarchical framework in which 6-*O* sulfation, primarily mediated by *Hs6st2*, establishes a permissive extracellular signaling niche required for effective remyelination.

Although individual HS6ST isoforms have been reported to exhibit differences in substrate specificity (Habuchi et al., 2000), biochemical analyses suggest largely overlapping catalytic activity among family members (Smeds et al., 2003), raising the possibility of functional redundancy. However, our data indicate that *Hs6st2* exerts a non-redundant role in the adult demyelinating CNS. Germline deletion of *Hs6st2* produces a selective reduction in specific 6-*O* sulfated disaccharides, particularly ΔUA-GlcNS(6S) (Moon et al., 2024), consistent with our observations in *Hs6st2* null OPCs. In fibroblasts derived from *Hs6st1/2* double knockout mice, 6-*O* sulfation is markedly reduced but not completely abolished, and growth factor responsiveness is selectively altered (Sugaya et al., 2008), illustrating that partial quantitative changes in 6-*O* sulfation can produce context-dependent signaling defects without global disruption of HS structure. More generally, mice lacking *Hs6st2* exhibit relatively mild alterations in hippocampal gene expression and behavior (Moon et al., 2024). In humans, HS6ST2 mutations are associated with intellectual disability, and neurodevelopmental disease (Paganini et al., 2019), as well as neuroticism (Luciano et al., 2021). Together, these findings suggest that the impaired recruitment and differentiation observed in *Hs6st2*-deficient mice arise from quantitative and/or spatial alterations in 6-*O* sulfation rather than the complete abolition of this modification.

In contrast to the requirement for appropriate 6-*O* sulfation, deletion of *Hs2st1* enhanced oligodendrocyte recruitment and differentiation, a phenotype that initially appears paradoxical. OPCs have previously been reported to exhibit relatively high levels of 2-*O* sulfation compared with mature oligodendrocytes (Properzi et al., 2008), suggesting that developmental stage-specific sulfation patterns may influence lineage progression. However, disaccharide profiling revealed a marked compensatory increase in 6-*O* sulfated disaccharides in *Hs2st1*-deficient OPCs, consistent with prior reports that loss of 2-*O* sulfation redistributes sulfation toward 6-*O* positions (Merry et al., 2001; Merry and Wilson, 2002). This shift in sulfation balance phenocopied the pro-remyelinating effects observed following sulfatase inhibition (Saraswat et al., 2021b), suggesting that increased 6-*O* sulfation underlies the enhanced repair response in *Hs2st1* cKO mice. Critically, concurrent deletion of *Hs6st1/2* abolished the pro-differentiative effects of *Hs2st1* loss, demonstrating that elevated 6-*O* sulfation is required for this phenotype. These genetic epistasis experiments establish that 2-*O* sulfation does not independently promote remyelination but instead constrains repair indirectly through modulation of 6-*O* sulfation levels. Notably, although *Hs2st1* deletion failed to rescue the deficits observed in 6-*O* deficient mice, combined loss of 2-*O* and 6-*O* sulfation further reduced the proportion of CC1⁺ oligodendrocytes at 7 dpl relative to 6-*O* deficiency alone. This suggests that 2-*O* sulfation-dependent signaling may contribute to differentiation in a context-dependent manner, perhaps via modulation of Slit/Robo-dependent signaling that exhibits selective 2-*O* sulfation requirements (Pratt et al., 2006; Conway et al., 2011), and that this effect emerges when dominant 6-*O* mediated pathways are disrupted. Together, these findings are consistent with a hierarchical model in which the relative balance between 2-*O* and 6-*O* sulfation, rather than absolute 2-*O* modification *per se*, determines the regenerative competence of OPCs within demyelinating lesions.

The reciprocal phenotypes observed following manipulation of sulfotransferases and sulfatases further underscore the central role of 6-*O* sulfation in remyelination. Deletion of *Hs6st2* reduces 6-*O* incorporation during HS biosynthesis and impairs OPC recruitment and differentiation, whereas loss of *Sulf1/2* preserves 6-*O* sulfation on extracellular HS chains and enhances repair. Rather than reflecting opposing mechanisms, these findings converge on a common principle that the level of 6-*O* sulfation is a key determinant of regenerative competence within demyelinating lesions. Under this model, *Hs6st2* establishes a permissive 6-*O* sulfated scaffold required for effective growth factor signaling, while sulfatases dynamically edit this modification to fine-tune pathway activity. Thus, both biosynthetic and post-synthetic regulation of 6-*O* sulfation operate along a shared regulatory axis, and shifts in 6-*O* balance, through reduced synthesis, increased retention, or compensatory redistribution following 2-*O* loss, directly influence remyelination outcomes.

Although 6-*O* sulfation emerged as the key HS modification regulating OPC repair, its relationship to canonical WNT/BMP signaling appears context-dependent. In *Hs2st1* cKO lesions, OPCs exhibited reduced expression of the WNT target *Apcdd1* and the BMP target *Id4*, consistent with attenuation of inhibitory transcriptional programs during the differentiation phase. We previously observed a similar reduction in *Apcdd1* and *Id4* following conditional deletion of *Sulf1/2* in OPCs or pharmacological inhibition of sulfatase activity with PI-88, both of which preserve or enhance 6-*O* sulfation within lesions (Saraswat et al., 2021b). Together, these findings suggest that elevated or retained 6-*O* sulfation is sufficient to dampen canonical WNT/BMP signaling in OPCs. In contrast, deletion of *Hs6st2* or combined loss of 6-*O* sulfotransferase activity did not significantly alter *Apcdd1* or *Id4* expression at 7 days post lesion, despite clear impairments in oligodendrocyte accumulation and maturation. These data suggest that although increased 6-O sulfation can suppress inhibitory WNT and BMP signaling, reduced 6-*O* sulfation does not further augment canonical pathway output at this stage. Instead, impaired remyelination under conditions of reduced 6-*O* sulfation likely reflects WNT- and BMP-independent mechanisms that regulate OPC recruitment and responsiveness to permissive cues.

Collectively, these data indicate that HS *O*-sulfation on OPCs regulates remyelination through both cell-intrinsic and potentially cell-extrinsic mechanisms. Conditional deletion of *Hs6st1*, *Hs6st2*, or *Sulf1/2* within OPCs altered oligodendroglial recruitment and differentiation without producing overt changes in astrocytic or microglial activation, supporting a predominantly OPC-intrinsic role for 6-*O* sulfation in controlling extracellular signaling within the oligodendrocyte lineage. In contrast, *Hs2st1* deletion was accompanied by reduced astrocytic reactivity and altered astrocyte gene expression, including an increased proportion of Timp1-expressing astrocytes, suggesting that 2-*O* sulfation may additionally influence lesion responses through OPC-dependent communication with astrocytes. Across all models, changes in oligodendrocyte lineage density occurred largely independently of major alterations in proliferation or apoptosis, consistent with sulfation-dependent regulation of OPC signaling competence and differentiation state. Although the mechanisms underlying these glial interactions remain unresolved, the divergent phenotypes observed following manipulation of 2-*O* and 6-*O* sulfation suggest that distinct HS sulfation patterns differentially coordinate both intrinsic oligodendrocyte responses and intercellular signaling within the lesion environment.

In summary, our findings identify HS *O*-sulfation as a tunable extracellular regulator of remyelination in which 6-*O* sulfation represents the dominant determinant of OPC regenerative competence. We demonstrate that *Hs6st2*-mediated 6-*O* incorporation establishes a permissive signaling environment required for effective recruitment and differentiation, whereas loss of 2-*O* sulfation enhances repair indirectly through compensatory elevation of 6-*O* sulfation. The convergence of genetic and pharmacological manipulation along this axis underscores the importance of maintaining an optimal level of 6-*O* sulfation within demyelinating lesions. Given that HS-modifying enzymes are dynamically regulated in both experimental models and human MS tissue, targeted modulation of the 6-*O* sulfation axis may represent a tractable extracellular strategy to enhance endogenous myelin repair. More broadly, these results suggest that precise regulation of ECM sulfation patterns constitutes a fundamental mechanism governing CNS plasticity and regeneration.

## Acknowledgements

This work was supported by the Department of Defense Congressionally Directed Medical Research Programs (HT9425-23-1-0682) and National Institute of Neurological Disorders and Stroke grant R01NS104021 to F.J.S., and by National Institute of Dental and Craniofacial Research grant R01DE031273 to D.X.

## Conflict of Interest

The authors declare no competing financial interests.

## Supplemental Figures

**Supplementary Figure 1:**
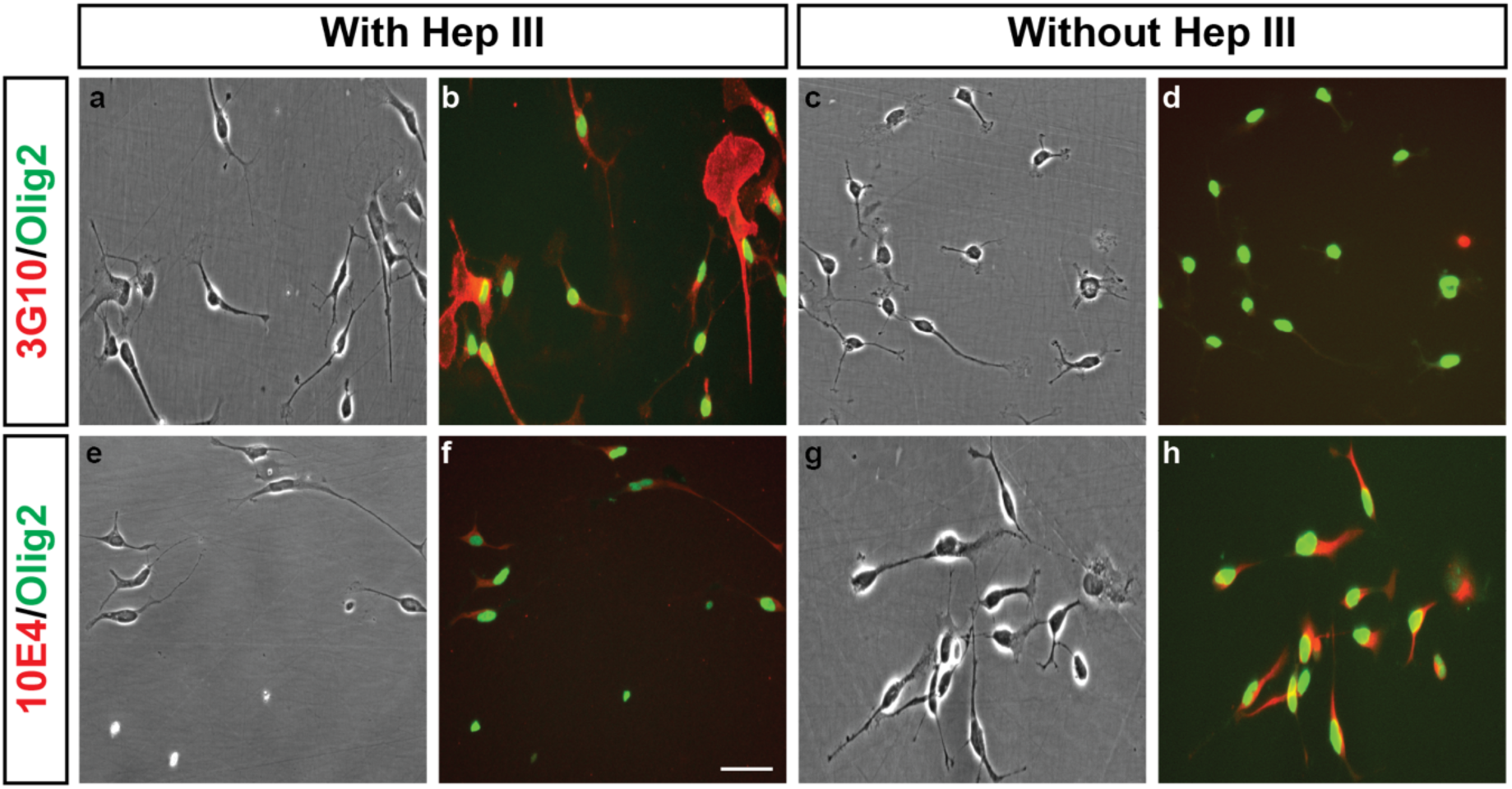
Expression of the neo-epitope of heparan sulfate (HS) and N-sulfated HS by human OPCs. The neo-epitopes of HS and N-sulfated HS were identified using 3G10 (red) and 10E4 (red), on Olig2 (green)-expressing human CD140a^+^ primary OPCs, isolated as previously described (Sim et al., 2011). HEP III treatment cleaves the HS chain from the core protein, thereby generating the 3G10 epitope; however, HEP III treatment shows no immunoreactivity with the 10E4 antibody, indicating a lack of binding to the cleaved structures. To validate the specificity of 3G10 and 10E4, the cells were treated with and without HEP III, as shown in panels (**b**, **f**) and (**d**, **h**), respectively. Phase contrast images of human primary OPCs are presented in panels **a**, **c**, **e**, and **g**, providing a clear view of cellular morphology. Scale: 50 µm.

**Supplementary Figure 2:**
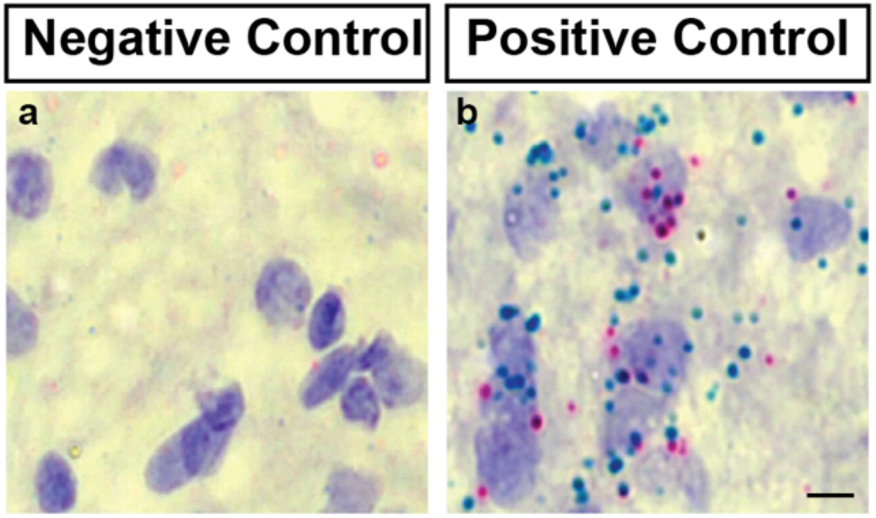
Negative and Positive controls of BaseScope assay. Validated the sensitivity and specificity of the 1ZZ BaseScope technology by assessing staining with the negative control probe (bacterial mRNA dapB; **a**) and the positive control probes (positive control: housekeeping gene PPIB and POLR2A; **b**). Scale: 50 μm.

**Supplementary Figure 3:**
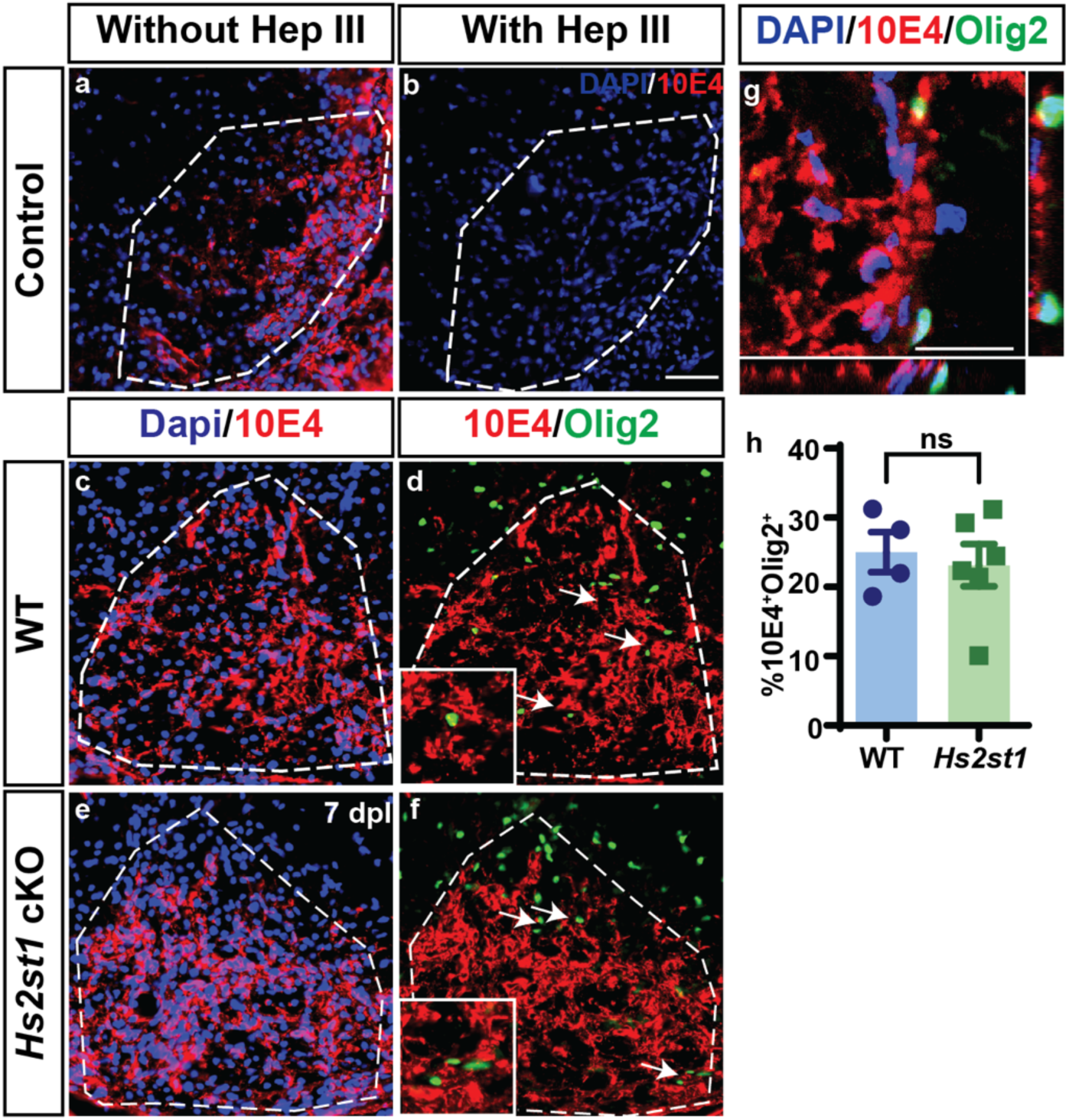
Conditional knockout of *Hs2st*1 in OPCs does not influence the overall expression level of HS in oligodendrocyte lineage cells. OPC-specific *Hs2st1* cKO was induced in adult *NG2CreERT2: Hs2st1^fl/fl^* mice by intraperitoneal administration of 200 mg/kg tamoxifen every other day (total of 5 injections). Cell surface HS expression in WT and *Hs2st1* cKO was assessed by immunostaining with the anti-HS monoclonal antibody (10E4) in adult mice following lysolecithin-induced demyelination at 7 dpl. **a**, HS chains were detected inside lesion (white dotted line) by 10E4 antibody (red) and were co-labeled with nuclei (blue) in the absence of HEP III. **b**, 10E4 staining was not detected after HEP III treatment. 10E4 expression on oligodendroglial lineage cells in the lesion was examined by co-labeling 10E4 (red), DAPI (blue) and Olig2 (green) in WT (**c, d**) and *Hs2st1* cKO (**e, f**). **g**, Orthogonal reconstruction shows 10E4 (red) colocalization with Olig2^+^ (green). **h,** Proportion of 10E4^+^Olig2^+^ among Olig2^+^ cells (n = 4-6 mice). *Hs2st1* cKO had no effect on N-sulfated HS chains. Mean ± SEM. Scale: **a**-**f**, 50 µm; **g**, 20 µm.

**Supplementary Figure 4:**
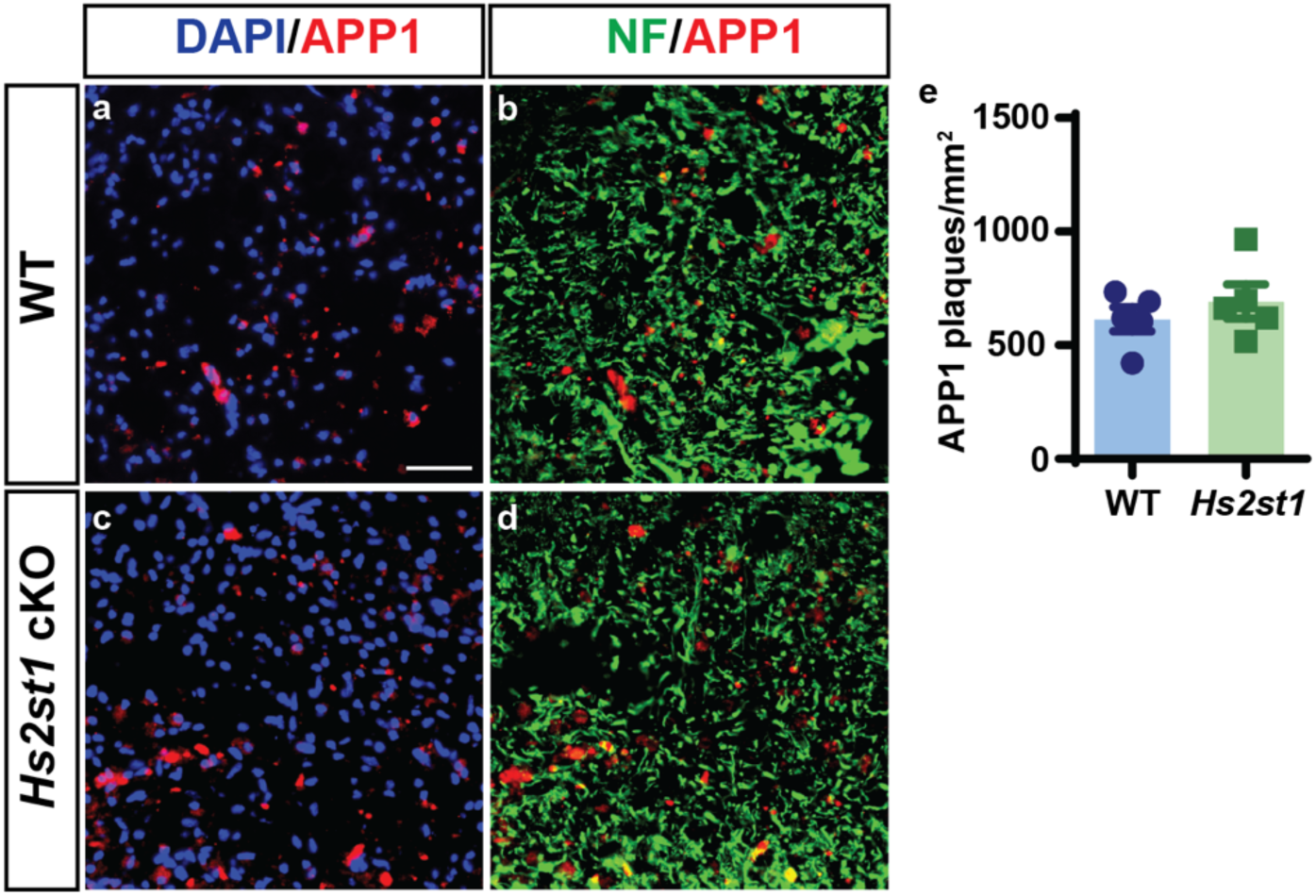
*Hs2st1* cKO in OPCs does not influence axonal injury following lysolecithin-induced demyelination. Tamoxifen was administered intraperitoneally at 200 mg/kg (5 injections, every other day) to induce *Hs2st1* cKO in the *NG2CreERT2: Hs2st1^fl/fl^* mice. After 7 days of the last tamoxifen injection, animals underwent intraspinal lysolecithin-induced demyelination. Axonal injury was assessed by APP1 immunofluorescence on neurofilament (NF)^+^ axons (heavy and light chains) in the WT (**a**, **b**) and *Hs2st1* cKO (**c**, **d**) at 5 dpl. The proportion of colocalized APP plaques on NF^+^ axons was quantified (**e**) in the WT and *Hs2st1* cKO. Mean ± SEM (n = 5 mice/group). Scale: 50 µm.

**Supplementary Figure 5:**
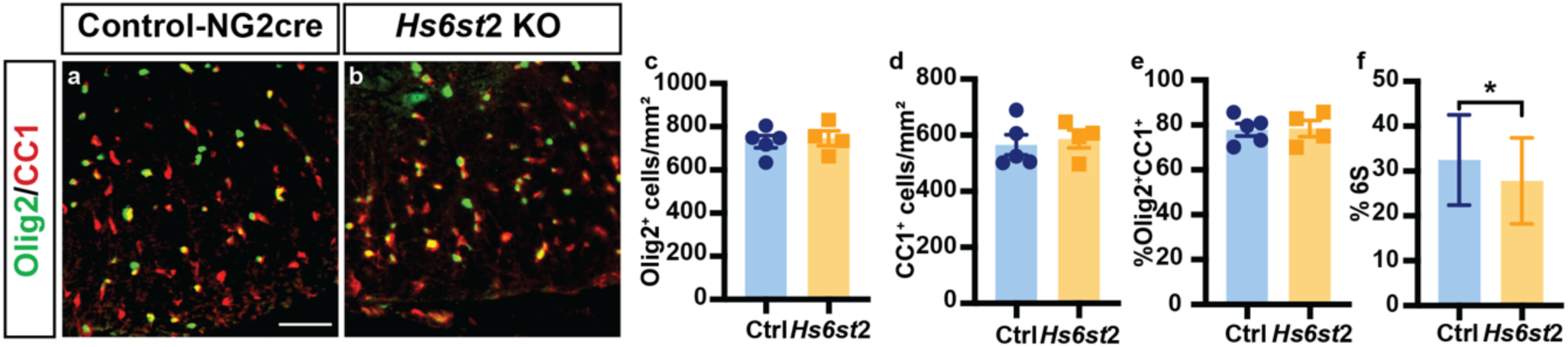
*Hs6st2* deletion does not affect OPC and oligodendrocyte density in the uninjured mouse spinal cord. The effect of germline *Hs6st2* deletion on oligodendrocytes in the adult spinal cord at 8-weeks was assessed by immunofluorescence. **a-b**, Oligodendroglial lineage cells were identified by Olig2 (green), and mature oligodendrocytes by CC1 (red). The Olig2^+^ and Olig2^+^CC1^+^ cell density (**c**, **d**) and percentage of Olig2^+^CC1^+^ cells among the Olig2^+^ cell population (**e**) was quantified in uninjured adult mouse spinal cord of control and *Hs6st2* KO mice (n = 5 and 4 for control and KO mice, respectively). **f**, The composition of HS disaccharides in OPCs isolated from WT and *Hs6st2* KO mice was assessed by LC-MS/MS. The molar fraction of 6S modified disaccharides was moderately decreased in *Hs6st2^−/−^*OPCs compared with OPCs isolated from control mice (paired Student’s t-test *p* < 0.05, n = 3 independent paired isolations per genotype). Mean ± SEM shown. Scale: 50 µm.

## Notes

### Competing Interest Statement

The authors have declared no competing interest.

